# A Novel Method for Across-Chromosome Phasing without Relative Data

**DOI:** 10.64898/2026.02.15.706064

**Authors:** Emmanuel Sapin, Kristen M. Kelly, Matthew C. Keller

## Abstract

**Motivation:** Across-chromosome phasing identifies which haplotypes of different chromosomes come from the same parent. This differs from within-chromosome phasing, which uses linkage disequilibrium patterns to determine which alleles were co-inherited within each chromosome but does not match haplotypes across different chromosomes. While across-chromosome phasing can be conducted using genotypes from parents or close relatives, current methods perform poorly for samples of unrelated individuals. Here, we introduce a novel approach for across-chromosome phasing that employs a window-based SNP-similarity metric, eliminating the need for data from close relatives or detection of identical-by-descent haplotypes.

**Results:** Using UK Biobank offspring with both parents genotyped as a gold standard, we evaluated the performance of our method by phasing the offspring without using parental data. In genomic data with no within-chromosomal phase errors, our algorithm achieved a mean across-chromosome phasing accuracy of 95%, with 53% of individuals phased perfectly. When data was pre-phased computationally using a standard within-chromosomal phasing algorithm, mean accuracy for across-chromosome phasing dropped to 83.1%. Thus, our method is limited primarily by the accuracy of within-chromosome phasing, and can approach near perfect across-chromosome phasing accuracy as within-chromosome phasing accuracy improves.

## 1 Introduction

As a diploid species, humans possess two copies of each autosomal chromosome, one inherited from each parent. Genetic data typically indicates which alleles are present at each locus, but not which alleles co-occur together on the same copy of a chromosome. Phasing—the process of separating diploid genotypes into two haploid sets of alleles (haplotypes) that reside on different copies of a chromosome—is a key challenge in genetics. The term “phasing” generally refers to within-chromosome phasing, where the process of assigning alleles to haplotypes occurs independently for each chromosome. Improving the quality of within-chromosome phasing is an area of significant interest [22, 35] due to its numerous applications in genetics, such as genotype imputation [7], local ancestry inference [8, 9], Mendelian disease research [23], and detection of haplotypes that are identical-by-descent (IBD) between individuals [10]. Tools such as Beagle [11], Eagle2 [20] and Shapeit2 [5] have improved the accuracy and computational efficiency of within-chromosome phasing considerably.

The rate of switch errors between two heterozygous sites, where the haplotype phasing algorithm incorrectly switches the assignment of alleles to a different haplotype, can be as low as 0.0016 [12] in well-phased data.

Across-chromosome phasing matches haplotypes on different chromosomes by their parent of origin after within-chromosome phasing has been completed. Accurate across-chromosome phasing reveals which chromosomal haplotypes were transmitted together from the same parent, although it does not necessarily specify which set of haplotypes is paternal or maternal in origin. This approach can enhance relatedness inference [38] and pedigree reconstruction [32, 33], support identification of each parent’s population of origin [34], enable assessment of parent-of-origin effects [13, 14], and boost power in linkage or genome-wide association studies (GWAS) by facilitating analysis of individuals not directly genotyped [21, 25]. Such augmented data also increases the power of GWAS by proxy (i.e., analyzing a child’s genomic data in place of a proband parent’s)[26, 27], as it differentiates half of each parent’s genome. Additionally, across-chromosome phased data can help to clarify changes over time in the strength of assortative mating, as correlations of polygenic scores between the (across-chromosome) phased haplotypes within a person reflect assortment patterns in the parental generation [15].

Both within- and across-chromosome phasing of genetic data is straightforward and almost perfectly accurate when genetic data from both parents is available for comparison. SNPs where the offspring is heterozygous and at least one parent is homozygous make it possible to determine which parent transmitted each allele, and best-guesses based on within-chromosome phasing of flanking SNPs can be used to fill in gaps where the offspring and both parents are heterozygous [36]. However, most datasets do not have genetic data for participants’ parents, and across-chromosome phasing without parental data is a more difficult problem with few established methods. Existing approaches often have requirements that are difficult to meet, such as the presence of other (non-parental) close relatives in the data (e.g., [21, 24]) or are intended for special circumstances such as interspecific hybrids or allopolyploid species [4].

Three recent approaches enable across-chromosome phasing in cohorts comprising individuals who are not necessarily closely related. In [13] the proposed method is predicated on the concept of surrogate parents. A surrogate parent is defined as a relative of the focal individual who is inferred to be related to one of the focal individual’s biological parents, with this relationship established through analysis of sex chromosome data. This approach demonstrates near-perfect performance; however, the reported results pertain only to a subset of the dataset consisting of individuals for whom surrogate parent relationships could be reliably identified.

In Noto et al. (2022) [17], locations are identified where the focal individual and other individuals share Identical By Descent (IBD) segments at least 10 centimorgans (cM) long on at least two different chromosomes. Haplotypes on different chromosomes that are mutually IBD between a pair of individuals were probably inherited from a common ancestor and so can be inferred as being in phase (inherited from the same parent). Clusters of overlapping segments are then created and matched to allow for across-chromosome phasing over the whole genome. The accuracy of this approach depends on the number of relatives present in the sample. In U.S.-based samples of predominantly European ancestry that are not enriched for close relatives, sample sizes of 10 million or more are required to yield sufficient numbers of both close and distant relatives to achieve error rates of approximately 5%. In the approach by Cole et al. [18], which was developed at the same time and in parallel with the approach introduced in the current manuscript, IBD segments of 5 cM or longer are detected for the focal individual, then a signed graph is built. Each IBD segment is a vertex of the graph, and signed edges are created between overlapping IBD segments to indicate whether the segments overlap on the same haplotype or on opposite haplotypes. Segments are grouped into tiles when they overlap by at least 3 cM. A clustering algorithm is then applied to the graph to create two clusters corresponding to tiles of segments inherited from each of the two parents. The approach in [18] is more robust to within-chromosome phasing errors in the data than the approach in [17], but both approaches depend on the ability to detect numerous IBD segments that meet minimum length requirements. These methods are therefore computationally intensive and work best in large datasets or datasets with close relatives to produce a sufficient number of long IBD segments.

This paper introduces a novel approach to across-chromosome phasing that leverages correlations of a SNP-based similarity measure across-chromosomes, eliminating the need for explicit IBD segment calling. It is effective in smaller datasets (fewer than 500,000 individuals) and performs well when individuals lack the close relationships necessary to share long IBD segments. We detail the methodology and validate its performance using a dataset that includes offspring from complete parent-child trios, enabling ground-truth comparisons to assess phasing accuracy. Without using data from close relatives, our method achieved a median accuracy of 85.93% (mean: 83.10%) when applied to data that had been within-chromosome phased using a standard algorithm. When the initial within-chromosome phasing was error-free, median accuracy increased to 100% (mean: 95%).

## 2 Overview of Approach

We developed and evaluated our method using data from the UK Biobank [19], focusing on individuals of European descent. This choice was driven by the method’s altered performance in mixed-ancestry samples and the fact that individuals of European descent were overwhelmingly the largest group in the UK Biobank. We used genetic principal components to identify the smallest 4-dimensional hypersphere containing all individuals with self-described white British ancestry, then selected all individuals within that space (regardless of self-described ancestry) as our eligible sample (435,187 individuals). Next, we selected biallelic SNPs that were present on both the UK BiLEVE and UK Biobank Axiom arrays, had a minor allele frequency (MAF) of at least 5%, and that had a Hardy-Weinberg equilibrium chi-square test p-value greater than 0.0005. This resulted in a final dataset of 330,005 SNPs.

We used data from 978 trio families, each consisting of an offspring and both parents, to evaluate phasing accuracy in the offspring. Using parental genotypes, we established a “gold standard” or ground truth for across-chromosome phasing, supplemented by within-chromosome phase information inferred with Shapeit2 [6], at sites flanking SNPs where all three family members were heterozygous. This gold-standard phasing is expected to be nearly 100% accurate, with errors arising only from SNP-calling inaccuracies or within-chromosome phase switch errors near mutually heterozygous sites. Such switch errors are typically span only one or two heterozygous SNPs, as they are corrected at the next site at which one family member is homozygous.

After establishing the ground truth for across-chromosome phasing in the trio offspring, we excluded their parents and any other first-degree relatives, and applied all steps of our phasing method using the offspring as focal individuals. The parameters used in the across-chromosome phasing procedure were selected to optimize phasing accuracy within this trio offspring sample. As discussed in the Results section, we do not believe this led to upwardly biased estimates of phasing accuracy due to overfitting, despite parameter tuning being performed on the same sample of 978 individuals.

All other “non-focal” individuals in the full European-ancestry sample were retained and used to phase the focal individuals across chromosomes. This design allowed us to evaluate phasing accuracy under conditions more representative of large-scale cohorts such as the UK Biobank, where most individuals lack first-degree relatives.

Nevertheless, we observed that levels of more distant relatedness were higher among trio offspring than in the broader UK Biobank sample. For example, 15.9% of trio offspring had at least one second-degree relative in the dataset (Figure 2, Table 1 in the Supplement), compared to only 4.1% among other individuals in the UK Biobank. To enable generalizable inferences across samples with varying levels of relatedness, we present phasing accuracy stratified by each individual’s closest degree of relatedness in the dataset.

**Table 1.** Correlations between 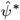 vectors across haplotypes of two windows for a focal individual.

| | | Window $w_h$ | |
| --- | --- | --- | --- |
| | | Hap $A$ | Hap $B$ |
| Window $w_g$ | Hap $A$ | $r(\hat{\psi}_{A,w_g}^*, \hat{\psi}_{A,w_h}^*)$ | $r(\hat{\psi}_{A,w_g}^*, \hat{\psi}_{B,w_h}^*)$ |
| | Hap $B$ | $r(\hat{\psi}_{B,w_g}^*, \hat{\psi}_{A,w_h}^*)$ | $r(\hat{\psi}_{B,w_g}^*, \hat{\psi}_{B,w_h}^*)$ |

All UK Biobank individuals were initially phased within chromosomes using Shapeit2 [19]. For trio families, focal individuals (offspring) were phased without incorporating parental or sibling genotypes, as described earlier. Because across-chromosome phasing depends on the accuracy of within-chromosome phasing, we applied a multi-stage error-correction algorithm to reduce genotyping and phase errors across the full sample. This method leverages haplotype sharing with long matching segments to identify inconsistencies and iteratively refine phase accuracy. Further details on phasing and correction procedures are provided in the Supplement.

Our method analyzes one focal individual at a time, using a SNP-based similarity metric to compare one haplotype of the focal individual to both haplotypes of every other individual in the sample within fixed, non-overlapping windows. For each haplotype of a non-focal individual, the higher of the two haplotypic SNP-similarity values for a given window is selected, ensuring that the most relevant haplotype pairing is used. This approach minimizes the impact of phasing errors in non-focal individuals, because the selection of which of the non-focal individual’s chromosomes is more similar for each window is based only on SNPs from within that window, allowing each window to match to either of the non-focal individual’s chromosomes. In particular, to quantify haplotypic SNP similarity between focal and non-focal individuals within each window, we compute the maximal SNP similarity for one haplotype (*A*) of the focal individual for a given window *w*_*g*_ on a given chromosome. This results in a vector 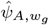 of length *n* − 1, where each element in the vector represents the SNP similarity between haplotype *A* of the focal individual and the most similar haplotype of each non-focal individual of window *w*_*g*_ on the same chromosome. Similarly, we calculate 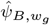 for the focal individual’s second haplotype (*B*) for window *w*_*g*_, producing another vector of *n* − 1 SNP similarity values. The labels *A* and *B* are assigned arbitrarily for each pair of chromosomes and do not yet reflect across-chromosome phase.

After calculating haplotypic similarity values for each window across the genome between the focal individual and all non-focal individuals, we examine the 2 × 2 correlation matrices of SNP similarity values between pairs of haplotypes at each chromosomal window. Specifically, we compute the correlation between the vectors 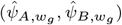 and 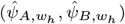, where *w*_*g*_ and *w*_*h*_ refer to windows *w*_*g*_ and *w*_*h*_ on either the same or different chromosomes, respectively. Correlations between these 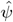 vectors are expected to be stronger when both haplotypes are inherited from the same parent (e.g., the maternal haplotype of window 1 on chromosome 4 and maternal haplotype of window 3 on chromosome 7) than when they are inherited from different parents (e.g., the maternal haplotypes of window 1 on chromosome 4 and the paternal haplotype of window 3 on chromosome 7). By iteratively pairing haplotypes across windows and chromosomes based on these 2×2 correlation matrices—beginning with the pair showing the strongest evidence of shared parental origin—we can cluster windows inherited from the same parent, enabling accurate across-chromosome phasing.

The underlying intuition is that a common ancestor of the focal and non-focal individuals may sometimes transmit two or more IBD segments across different chromosomes. In the absence of inbreeding, such segments should be inherited from either one parent of the focal individual or the other, depending on which side of the family tree the common ancestor is from, meaning they should be in across-chromosome phase with one another. Absent inbreeding, focal haplotypes that show elevated SNP similarity values with the same non-focal individuals across two or more chromosomal windows are likely inherited from the same parent. This pattern should lead to inflated correlations of 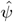 between haplotypic windows that are in phase with one another while avoiding the need to identify IBD segments.

While 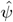 is typically influenced by IBD segments inherited from a common ancestor, it remains effective when IBD segments are too short to formally detect. The 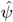 metric is also informative when the focal individual's parents belong to two different ancestral sub-populations. In such cases, differences in allele frequencies due to population stratification, including fine-scale stratification [37], can affect 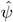 values. Individuals from the same sub-population as one of the focal individual's parents will exhibit higher 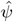 values for haplotypes originating from that parent. This creates a similar pattern of correlation between the lists of 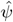 values and allows population stratification in the parental generation to assist in across-chromosome phasing.

The implementation was executed within a multi-node computational environment [2], employing parallelization techniques in the C programming language. The source code has been made publicly accessible online [1], thereby facilitating reproducibility of the results for researchers with authorized access to the UK Biobank dataset.

The remainder of this manuscript describes the details of our across-chromosomal phasing approach and its performance. Section 3 describes the calculation and use of the 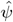 metric and our algorithm to use this metric for across-chromosome phasing. Section 4 evaluates the accuracy of the across-chromosomal phasing using the 978 trio offspring, The last section summarizes the paper.

## 3 Across-chromosome phasing algorithm

### 3.1 Calculation of the 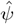 metric

For the purposes of this study, we computed the 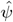 metric for each focal individual using only non-focal individuals whose genome-wide SNP relatedness (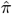[3]) with the focal individual was less than 0.33. This threshold excludes parents, siblings, and offspring, ensuring that phasing accuracy estimates based on trio offspring are more representative of performance in population-based samples such as the UK Biobank. Note, however, that while close relatives such as parents or half-siblings could enable near-perfect phasing, full siblings are not informative for across-chromosome phasing in our framework, as they inherit both haplotypes from the same parents.

The formula for 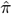 [3], a widely used measure of (diploid) SNP similarity [3], is shown in Equation 1, where *f* denotes the focal individual, *i* indexes non-focal individuals, *p*_*k*_ is the MAF for SNP *k, x*_*k*_ represents each individual’s diploid genotype at SNP *k*, and *m* is the total number of SNPs included in the calculation.

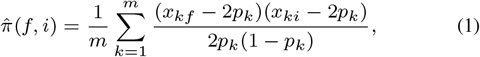

We modify the original formula to estimate similarity between haploid, rather than diploid, genotypes. A haploid version of the SNP similarity metric is given in Equation 2, where *A*_*f*_ and *A*_*i*_ represent haplotypes *A* from the focal and non-focal individuals, respectively. The variable *x*_*k*_ denotes the allele at SNP *k* within a sequence *s* located in window *w*_*g*_ on a given chromosome. The sequence *s* is defined as the longest contiguous run of SNPs for which the two haplotypes are identical; that is, 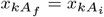 for all *k* ∈ *s* ⊆ *w*_*g*_. This construction ensures that 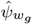 is always non-negative. The calculation of 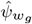 includes only SNPs that are heterozygous in the focal individual, as these are the only phase-informative sites. Each term in the sum is exponentiated by 1*/*5 to reduce the disproportionate influence of SNPs with low minor allele frequency (MAF). While rare SNPs still contribute more to the similarity score than common SNPs, the exponentiation dampens their impact, reducing noise in the resulting similarity metric. Unlike 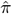, which is an average SNP similarity (the sum divided by the number of SNPs), 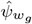 is a sum, allowing longer stretches of haplotypic identity to have greater influence.

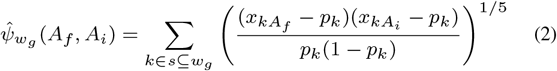

The length and number of windows within each chromosome are important parameters for accurate across-chromosome phasing. Windows that are too long are more likely to span phase errors, potentially propagating those errors to other regions. Conversely, windows that are too short may not contain enough SNPs to support reliable phasing. To define windows that are approximately independent, we identified the top 10 recombination hotspots on each chromosome—those with the highest ratio of genetic distance (cM) to physical distance (Mb)—using the deCODE sex-averaged recombination map (aau1043_datas3.gz) from [41]. These hotspots were used as boundaries, with the additional constraint that each window must span at least 25 cM. As a result, longer chromosomes had more windows than shorter ones; for example, chromosome 1 was divided into 5 windows, whereas chromosome 21 had 2. In total, we defined 78 windows genome-wide, with an average length of 44.08 cM (standard deviation = 17.38 cM) and a range from 19.3 cM to 102.4 cM. On average, each window contained approximately 4,231 SNPs. A map of window boundaries by chromosome is provided in the Supplementary Materials (Figures 6 to 27).

Two haplotypic-haplotypic similarity values are computed for each window: 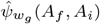 and 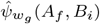, representing the similarity between the focal individual’s haplotype *Af* and the two haplotypes *Ai* and Bi of a non-focal individual. We then define 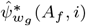 as the larger of these two similarity values, raised to the fourth power, as shown in Equation 3:

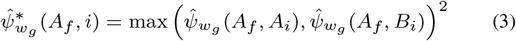

Exponentiating by 2 amplifies the contrast between low values of 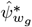—likely due to random similarity—and high values that are more indicative of IBD segments or shared ancestry. This metric serves as an estimate of the maximum haplotypic similarity between a given window *w*_*g*_ in haplotype *A*_*f*_ of the focal individual and the corresponding window in the two haplotypes of the non-focal individual.

Similarly, we compute 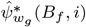 for the focal individual’s haplotype Bf using Equation 4:

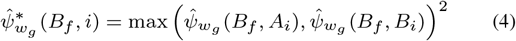

This approach ensures that, for each non-focal individual, only the haplotype most similar to the focal individual’s haplotype is considered. As a result, the method is more robust to within-chromosome phasing errors in non-focal individuals, since windows do not need to consistently align to the same haplotype. If a switch error in a non-focal individual spans multiple windows, it will only reduce 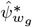 in the windows where the switch occurs, without affecting windows upstream or downstream of the window where the switch occurred. However, switch errors in the focal individual will still degrade across-chromosome phasing accuracy.

### 3.2 Usage of the 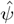 metric to phase across-chromosomes

For a given focal individual *f*, we define the vectors of length *n* − 1 whose elements are the values defined in Equations 3 and 4 as 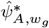 and 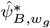 respectively, representing the maximum haplotypic similarity of window *w*_*g*_ between the focal individual’s haplotype A (or B) and the haplotypes of each of the non-focal individuals. These two vectors are computed for all 78 windows across all 22 autosomal chromosomes for a given focal individual. Our method for across-chromosomal phasing is based on these 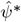 vectors. To phase two windows, *w*_*g*_ and *w*_*h*_, (which can exist on the same or, more often, on different chromosomes), we calculate correlations between the pair of vectors for each chromosome as shown in Table 1, where *r*(*x, y*) denotes the Pearson correlation between *x* and *y*.

Higher correlations along the diagonal of this matrix (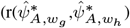) and 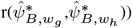 than the off-diagonal 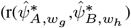 and 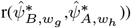 suggest that haplotype *A* of window *w*_*h*_ originates from the same parent as haplotype *A* of window *w*_*g*_, and similarly for haplotypes *B*. On the other hand, higher correlations along the off-diagonal of this matrix than the diagonal suggest the opposite: that haplotype *A* of window *w*_*h*_ pairs with haplotype *B* of window *w*_*g*_ and haplotype *B* of window *w*_*h*_ pairs with haplotype *A* of window wg.

In cases where two allelic windows (*A* and *B*), located on the same chromosome, both exhibit a high degree of correlation with a single haplotype (*A* or *B*) in another genomic window, the analytical procedure entails summing the correlations corresponding to identical haplotypes (i.e., *AA* and *BB*). In contrast, correlations associated with opposing haplotypes (*AB* and *BA*) are subtracted from this sum. This computation produces a unified quantitative measure, *λ*, that characterizes the relationship between a given pair of windows—whether situated on the same chromosome or on different chromosomes—as formally defined in Equation 5.

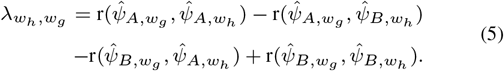

The sign of 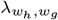 predicts whether haplotype *A* of window *w*_*h*_ and haplotype *A* of window *w*_*g*_ originate from the same parent (positive sign) or different parents (negative sign). In cases where parental genotypes are available, 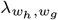 can be compared to the ground truth value to determine the accuracy of the across-chromosomal phasing.

### 3.3 Haplotype Matching Across Chromosomes

Our algorithm for matching haplotypes across windows—and thus across chromosomes—builds on the concepts introduced in the preceding sections. For each focal individual, the algorithm evaluates pairs of windows, denoted *w*_*g*_ and *w*_*h*_, which may lie on the same or different chromosomes. To account for the observation that switch errors between two windows occur with a frequency of less than half: if 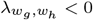 and the windows are located on the same chromosome, the value is scaled by a factor of 0.75. The algorithm then selects the pair that maximizes the absolute value of this adjusted 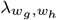 across all possible combinations. Subsequently, *w*_*g*_ and *w*_*h*_ are phased by integrating their haplotypes according to the sign of the selected *λ*. When 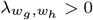, haplotype *A* of the first window is merged with haplotype *A* of the second (and *B* with *B*). Conversely, when 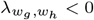, haplotype *A* is merged with haplotype *B* (and vice versa), thereby preserving the structural integrity of the haplotypic configuration.

Our algorithm for matching haplotypes across windows—and thus across chromosomes—builds on the concepts introduced in the preceding sections. For each focal individual, the algorithm begins by selecting a pair of windows, denoted *w*_*g*_ and *w*_*h*_, which may lie on the same or different chromosomes (and may or may not be adjacent if on the same chromosome). This pair is chosen to maximize the absolute value of 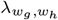 across all possible window combinations. The algorithm subsequently performs phasing of *w*_*g*_ and *w*_*h*_ across the chromosomal sequences, integrating their respective haplotypes in accordance with the sign of 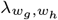. When 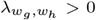, haplotype *A* from one genomic window is merged with the corresponding haplotype *A* of the other window, and the haplotype *B*s are similarly merged. Conversely, when 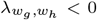, haplotype A of one window is merged with haplotype *B* of the other window (and vice versa), thereby preserving the structural integrity of the haplotypic configuration.

Next, a third window *w*_*l*_ is selected by again maximizing the absolute value of the correlation—now between *w*_*l*_ and the already merged haplotypes from *w*_*g*_ and *w*_*h*_, denoted *w*_*g,h*_. The haplotypes of *w*_*l*_ are aligned with those of *w*_*g,h*_, and the three windows are then merged into a single phased segment.

This iterative procedure continues, with each subsequent window selected and phased relative to the merged haplotypes of all previously processed windows. The process terminates once all windows across all chromosomes have been incorporated, yielding a fully across-chromosome phased haplotype for the focal individual. A formal description of the algorithm is provided in Supplement 2.

## 4 Results

We present the results of applying our across-chromosome phasing algorithm to UK Biobank data. To assess phasing performance, we define the *across-chromosome phasing accuracy* (*ACPA*) score as the proportion of SNPs that are phased together in the focal individual and correctly traced to the same biological parent, as illustrated in Figure 1. For SNPs at which the offspring and both parents are heterozygous, one allele is arbitrarily labeled as originating from Parent 1 and the other from Parent 2. If all alleles assigned to a given parent are in fact inherited from that parent, the *ACPA* score is 100%. An *ACPA* score of 50% is the expectation under random assignment, where alleles attributed to one parent actually originate equally from both parents.

**Figure 1.**
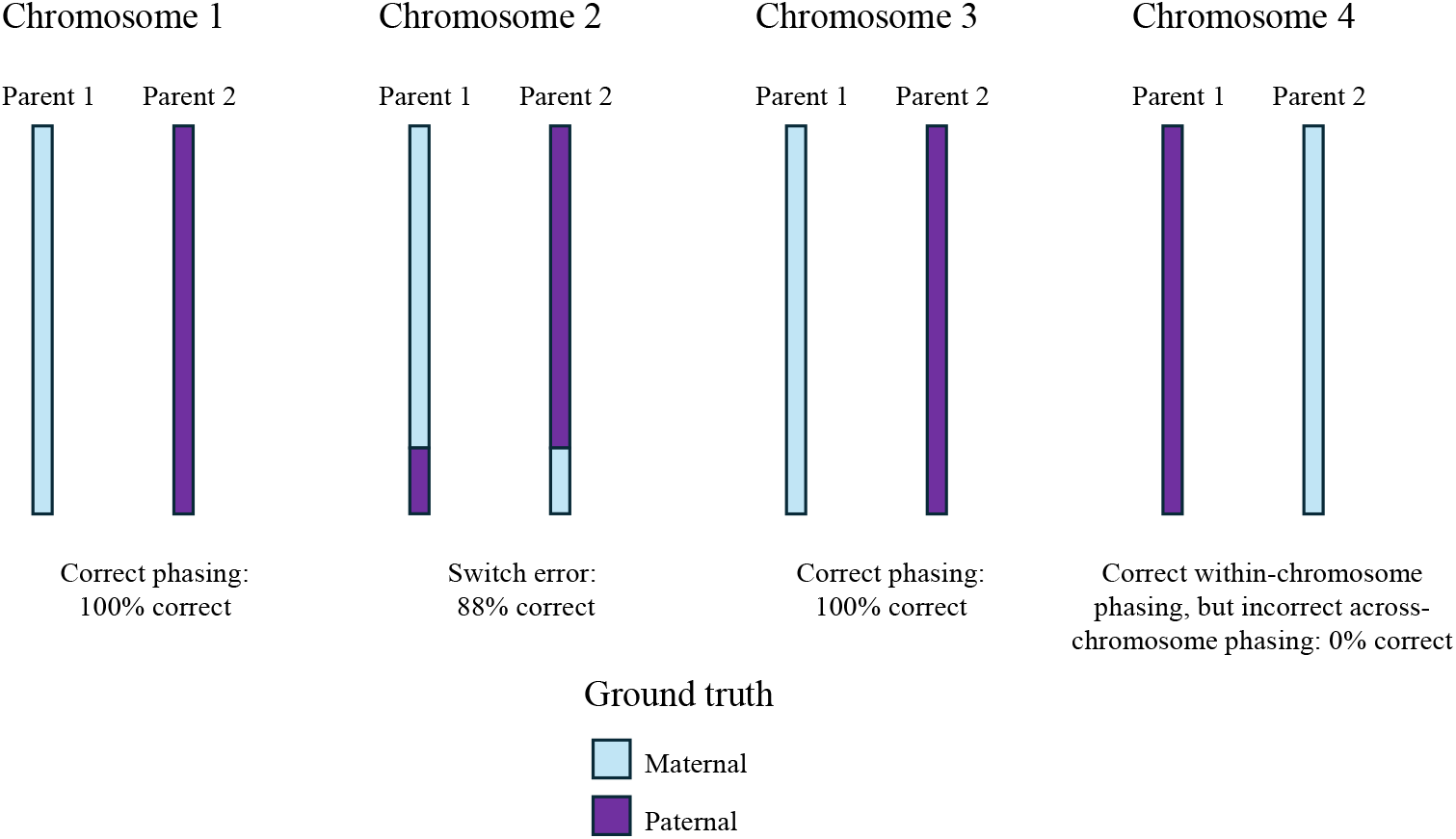
Simple example of across-chromosome phasing on four chromosomes of the same size, resulting in an *ACPA* score of (100+88+100+0)/4= 72%.

We first evaluated the accuracy of our algorithm using focal individuals (trio offspring) who had virtually no within-chromosome phasing errors (because parental genotypes were used to correct within-chromosome phasing). We then evaluated accuracy on data that included within-chromosome phasing errors (from being phased using Shapeit2 without first degree relative information used) as shown in the top of Figure 2. We present in the appendix results on the impact of the number of samples in the data (Section 5) as well as the quality of within chromosome phasing (Section 6).

**Figure 2.**
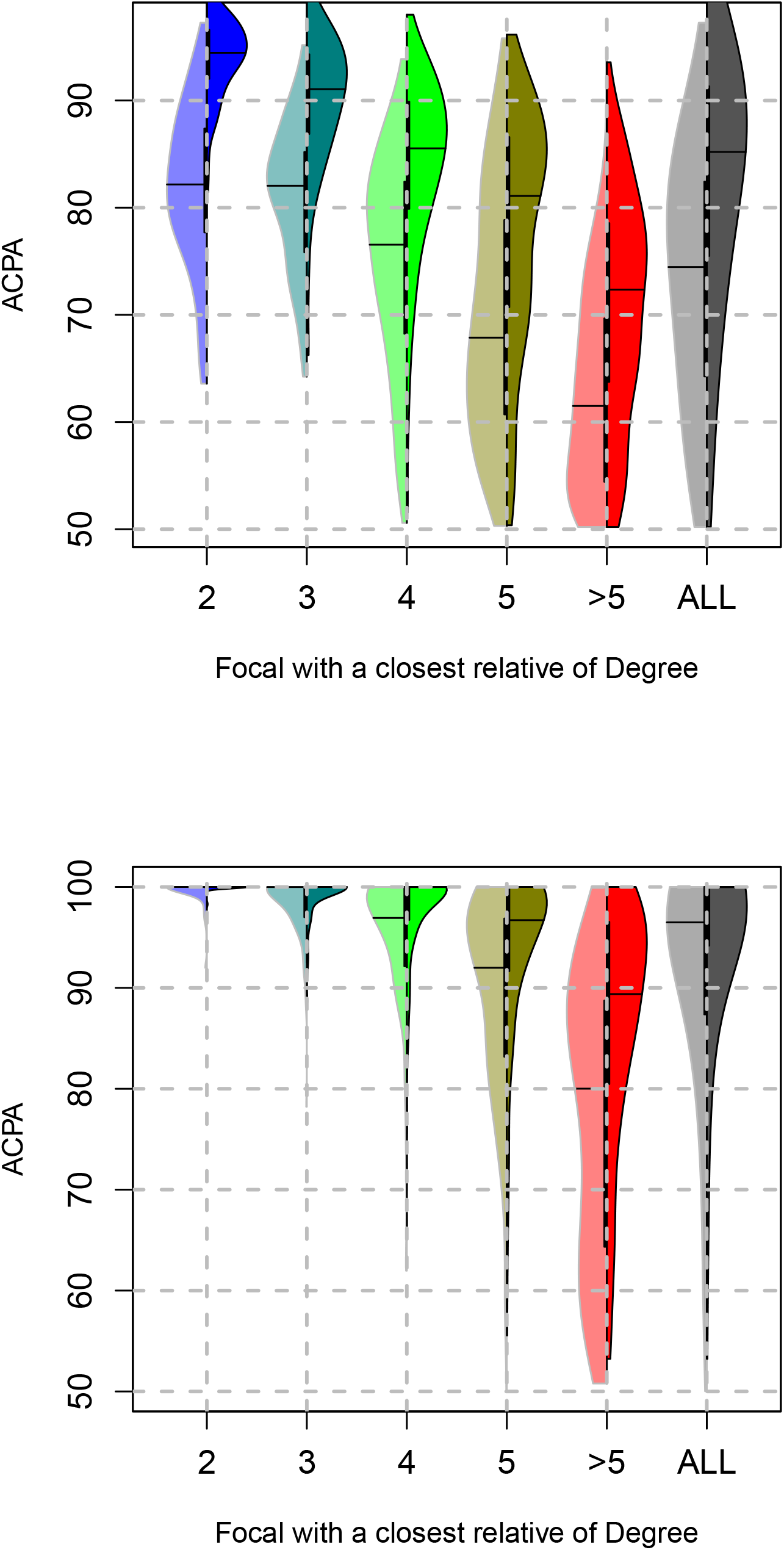
Violin plots of *ACPA* scores for trio offspring. The left half of each plot shows the *ACPA* scores for trio offspring when using the method of Noto et. al. while the right part of each violin shows *ACPA* scores using our method. The bottom plot shows results when input data has no within-chromosome phasing errors, while the top plot shows results when within-chromosome phasing errors are present. For each focal individual, color indicates the highest degree of relatedness (based on 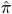) to any non-focal individual.

When no errors were present in the within-chromosome phasing, the median *ACPA* score across all offspring was 100%, with a mean of 95.3%. In contrast, when within-chromosome phasing errors were included, the median *ACPA* score dropped to 85.9%, with a mean of 83.10% (complete results are provided in Table 2). These results indicate that the primary limitation on the accuracy of across-chromosome phasing using our method is the quality of the initial within-chromosome phasing. Nonetheless, regardless of whether within-chromosome phasing errors were present, the presence of a close relative in the dataset consistently improved average phasing accuracy (Figure 2).

To benchmark our approach against existing methods, we applied the algorithm developed by Noto et al. [17] using their recommended IBD segment length threshold of at least 10 centimorgans. This method requires multiple non-focal individuals to share IBD segments with the focal individual on at least two chromosomes. When this requirement is not met—i.e., when some chromosomes lack sufficient IBD information—the method assigns phase at random for the affected chromosomes.

The results for the Noto method are shown in the left half of each violin plot in Figure 2, while results for our method are shown on the right. Across all comparisons, our approach outperformed the Noto method, both in individuals with and without close relatives, as reflected in the mean and median *ACPA* scores for trio offspring and parent–offspring (P/O) pairs following either computational (Shapeit2) or error-free phasing (Table 2).

To assess potential overfitting, we applied our method and the Noto algorithm to an independent sample of 3,718 offspring from parent–offspring (P/O) pairs of European descent who were not used in parameter tuning. Across-chromosome phasing accuracy is expected to be slightly lower in P/O than in trio offspring because (a) trio offspring in the UK Biobank happen to have more distant relatives in the dataset, providing stronger haplotypic similarity signal for phasing; and (b) trio data allow more complete and precise construction of ground truth for estimating accuracy. (It is important to re-emphasize that the trio accuracy estimates were not advantaged by parental information during within-chromosome phasing, and the same condition applied to the P/O analyses: parents were excluded during pre-phasing in both cases). In this validation sample, the median *ACPA* was 84.57%, with a mean of 81.73%, representing only a ∼1% decrease relative to trio offspring. In contrast, the Noto method showed a larger decline, with mean *ACPA* dropping from 73.3% in trio offspring to 68.1% in P/O offspring (Table **??**). This pronounced difference in performance decline indicates that our method’s accuracy in trio offspring is not inflated by overfitting. Additional P/O results are shown in the Supplement.

Finally, our method slightly outperforms the across-chromosome phasing results reported by Cole et al. [18]. In the supplementary materials of that study, the median *ACPA* score is 83.4% for 998 individuals from trio data self-described as White British or Irish (and parents with 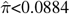) in the UK Biobank. When applying identical cohort selection criteria, the present approach attains a median *ACPA* score of 85.66% on those same 998 individuals.

## 5 Conclusion

Our approach—based on correlations of 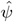 metrics—provides a flexible and effective framework for across-chromosome phasing, particularly in large cohorts of unrelated individuals. We introduce a novel method specifically designed for datasets in which individuals share few or undetectable IBD segments. This method leverages a new haplotypic similarity measure computed between pairs of genotyped individuals using computationally phased data. By comparing the haplotypic similarity profiles of a focal individual with those of all others in the dataset, we generate lists of similarity values that contain informative structure for inferring across-chromosome phase. Correlations between these lists across windows on separate chromosomes enable the alignment of haplotypes across chromosomes.

Applied to real data, our method outperforms existing IBD-based ap-proaches—such as that of Noto et al. [17]—particularly in individuals without close relatives. Moreover, it achieves high accuracy in a dataset of approximately 500,000 individuals, substantially smaller than the 10 million individuals recommended for the Noto method, highlighting the scalability and practical utility of our approach. The method is demonstrated to work well on human dataset and can be applied to any other species for which dataset of comparable sizes are available.

The results presented here are promising, but there are several ways in which they could be further improved. We have shown that our approach performs almost perfectly when the pre-existing phased data contains no within-chromosome phasing errors. Therefore, reducing the frequency of such errors would significantly enhance overall performance. Additionally, the method itself could be refined in several ways. We propose three potential improvements. First, our current approach treats windows independently, even when they lie on the same chromosome. However, within-chromosome phase information may be informative, as contiguous windows that are estimated to be in phase are more likely to reflect consistent across-chromosome phase. This information could be incorporated by introducing a loss function that penalizes phase configurations implying switch errors within chromosomes. Second, the integration of explicit information from close relatives could further improve phasing accuracy. For example, all genomic segments shared between an avuncular relative (e.g., an aunt or uncle) and another individual must originate from the same grandparental lineage—either maternal or paternal—providing a strong constraint on phase orientation. Third, across-chromosome phasing results could be used to refine within-chromosome phasing. Our method could be applied recursively, using improved across-chromosome phase estimates to inform a new round of within-chromosome phasing—potentially with smaller windows—thereby iteratively improving both within- and across-chromosome phase accuracy. Although the algorithm is conceptually general and could, in principle, be applied to other diploid species, practical performance will depend on factors such as adequate sample size and low levels of inbreeding, which are necessary to avoid parent-of-origin ambiguity. These potential enhancements underscore the flexibility and extensibility of our framework and suggest promising directions for future methodological development.

## References

[1] https://github.com/emmanuelsapin/AcrossChromosomesPhasing

[2] https://www.colorado.edu/rc/resources/blanca

[3] https://zzz.bwh.harvard.edu/plink/ibdibs.shtml

[4] Jia, K.-H., Wang, Z.-X., Wang, L., Li, G.-Y., Zhang, W., Wang, X.-L., Xu, F.-J., Jiao, S.-Q., Zhou, S.-S., Liu, H., Ma, Y., Bi, G., Zhao, W., El-Kassaby, Y.A., Porth, I., Li, G., Zhang, R.-G. and Mao, J.-F. (2022), SubPhaser: a robust allopolyploid subgenome phasing method based on subgenome-specific k-mers. New Phytol, 235: 801–809.

[5] Delaneau O, Zagury JF, Robinson MR, Marchini JL, Dermitzakis ET. Accurate, scalable and integrative haplotype estimation. Nat Commun. 2019 Nov 28;10(1):5436. doi: 10.1038/s41467-019-13225-y. PMID: 31780650; PMCID: PMC6882857.

[6] https://mathgen.stats.ox.ac.uk/genetics_software/shapeit/shapeit.html

[7] Marchini, J. and Howie, B. (2010). Genotype imputation for genome-wide association studies. Nature Reviews Genetics, 11(7):499–511.

[8] Maples, B., Gravel, S., Kenny, E., and Bustamante, C. (2013). RFMix: A Discriminative Modeling Approach for Rapid and Robust Local-Ancestry Inference. The American Journal of Human Genetics, 93(2):278–288.

[9] Freyman, W. A., McManus, K. F., Shringarpure, S. S., Jewett, E. M., Bryc, K., The 23 and Me Research Team, and Auton, A. (2021). Fast and Robust Identity-by-Descent Inference with the Templated Positional Burrows–Wheeler Transform. Molecular Biology and Evolution, 38(5):2131–2151.

[10] Zhou, Y., Browning, S. R., and Browning, B. L. (2020). A Fast and Simple Method for Detecting Identity-by-Descent Segments in Large-Scale Data. The American Journal of Human Genetics, 106(4):426–437.

[11] Browning, B. L., Tian, X., Zhou, Y., and Browning, S. R. (2021). Fast two-stage phasing of large-scale sequence data. The American Journal of Human Genetics, 108(10):1880–1890.

[12] Brian L. Browning, Sharon R. Browning, Statistical phasing of 150,119 sequenced genomes in the UK Biobank, The American Journal of Human Genetics, Volume 110, Issue 1, 2023, Pages 161–165

[13] Hofmeister, R.J., Cavinato, T., Karimi, R. et al. Parent-of-origin effects on complex traits in up to 236,781 individuals. Nature 646, 647–656 (2025). 10.1038/s41586-025-09357-5

[14] Hofmeister, R. J., Ribeiro, D. M., Rubinacci, S., and Delaneau, O. (2023). Accurate rare variant phasing of whole-genome and whole-exome sequencing data in the UK Biobank. Nature Genetics, 55(7):1243–1249.

[15] “Balbona, J.V., Kim, Y. and Keller, M.C. Estimation of Parental Effects Using Polygenic Scores. Behav Genet 51, 264–278 (2021). 10.1007/s10519-020-10032-w

[16] Ying Qiao, Jens G. Sannerud, Sayantani Basu-Roy, Caroline Hayward, Amy L. Williams, Distinguishing pedigree relationships via multi-way identity by descent sharing and sex-specific genetic maps, The American Journal of Human Genetics, Volume 108, Issue 1, 2021, Pages 68–83

[17] Noto, K., Ruiz, L. Accurate genome-wide phasing from IBD data. BMC Bioinformatics 23, 502 (2022). 10.1186/s12859-022-05066-2

[18] Cole M. Williams, Jared O’Connell, William A. Freyman, 23andMe Research Team, Christopher R. Gignoux, Sohini Ramachandran, Amy L. Williams. Phasing millions of samples achieves near perfect accuracy, enabling parent-of-origin classification of variants. bioRxiv 2024.05.06.592816; doi: 10.1101/2024.05.06.592816

[19] Bycroft, C. et al. (2018). The UK Biobank resource with deep phenotyping and genomic data. Nature 562, 203–209. 10.1038/s41586-018-0579-z

[20] Loh P-R, Danecek P, Palamara PF, Fuchsberger C, Reshef YA, Finucane HK, Schoenherr S, Forer L, McCarthy S, Abecasis GR, Durbin R, and Price AL. Reference-based phasing using the Haplotype Reference Consortium panel. Nature Genetics (2016).

[21] Kong, A., Masson, G., Frigge, M. L., Gylfason, A., Zusmanovich, P., Thorleifsson, G., Olason, P. I., Ingason, A., Steinberg, S., Rafnar, T., Sulem, P., Mouy, M., Jonsson, F., Thorsteinsdottir, U., Gudbjartsson, D. F., Stefansson, H., and Stefansson, K. (2008). Detection of sharing by descent, long-range phasing and haplotype imputation. Nature Genetics, 40(9):1068–1075.

[22] Sina Majidian, Fritz J Sedlazeck, PhaseME: Automatic rapid assessment of phasing quality and phasing improvement, GigaScience, Volume 9, Issue 7, July 2020, giaa078, 10.1093/gigascience/giaa078

[23] Beck CR, Carvalho CMB, Akdemir ZC, et al. Megabase length hypermutation accompanies human structural variation at 17p11.2. Cell. 2019;176 :1310–24.e10.

[24] Qiao Y, Jewett EM, McManus KF, Freyman WA, Curran JE, Williams-Blangero S, Blangero J; 23andMe Research Team; Williams AL. Reconstructing parent genomes using siblings and other relatives. bioRxiv [Preprint]. 2024 May 14:2024.05.10.593578.

[25] Lindholm Eva, Åberg Karolina, Ekholm Birgit, Pettersson Ulf, Adolfsson Rolf, and Jazin Elena E.. Reconstruction of ancestral haplotypes in a 12-generation schizophrenia pedigree. Psychiatric Genetics, 14(1), 2004.

[26] Liu Jimmy Z., Erlich Yaniv, and Pickrell Joseph K.. Case–control association mapping by proxy using family history of disease. Nature Genetics, 49:325–331, Jan 2017

[27] Hujoel Margaux L.A., Gazal Steven, Loh Po-Ru, Patterson Nick, and Price Alkes L.. Liability threshold modeling of case–control status and family history of disease increases association power. Nature Genetics, 52(5):541–547, May 2020.

[28] Kong Augustine, Thorleifsson Gudmar, Frigge Michael L., Vilhjalmsson Bjarni J., Young Alexander I., Thorgeirsson Thorgeir E., Benonisdottir Stefania, Oddsson Asmundur, Halldorsson Bjarni V., Masson Gisli, Gudbjartsson Daniel F., Helgason Agnar, Bjornsdottir Gyda, Thorsteinsdottir Unnur, and Stefansson Kari. The nature of nurture: Effects of parental genotypes. Science, 359(6374):424–428, 2018.

[29] Hwang Liang-Dar, Tubbs Justin D., Luong Justin, Lundberg Mischa, Moen Gunn-Helen, Wang Geng, Warrington Nicole M., Sham Pak C., Cuellar-Partida Gabriel, and Evans David M.. Estimating indirect parental genetic effects on offspring phenotypes using virtual parental genotypes derived from sibling and half sibling pairs. PLOS Genetics, 16(10):1–29, October 2020.

[30] Young Alexander I., Nehzati Seyed Moeen, Benonisdottir Stefania, Okbay Aysu, Jayashankar Hariharan, Lee Chanwook, Cesarini David, Benjamin Daniel J., Turley Patrick, and Kong Augustine. Mendelian imputation of parental genotypes improves estimates of direct genetic effects. Nature Genetics, 54(6):897–905, Jun 2022.

[31] Ramstetter Monica D., Shenoy Sushila A., Dyer Thomas D., Lehman Donna M., Curran Joanne E., Duggirala Ravindranath, Blangero John, Mezey Jason G., and Williams Amy L.. Inferring identical-by-descent sharing of sample ancestors promotes high-resolution relative detection. The American Journal of Human Genetics, 103(1):30–44, 2018.

[32] Staples Jeffrey, Qiao Dandi, Cho Michael H., Silverman Edwin K., Nickerson Deborah A., and Below Jennifer E.. PRIMUS: rapid reconstruction of pedigrees from genome-wide estimates of identity by descent. The American Journal of Human Genetics, 95(5):553–564, Nov 2014.

[33] Jewett Ethan M., McManus Kimberly F., Freyman William A., and Auton Adam. Bonsai: An efficient method for inferring large human pedigrees from genotype data. The American Journal of Human Genetics, 108(11):2052–2070, Nov 2021.

[34] Jagadeesan Anuradha, Gunnarsdóttir Ellen D., Ebenesersdóttir S. Sunna, Guðmundsdóttir Valdis B., Thordardottir Elisabet Linda, Einarsdóttir Margrét S., Jónsson Hákon, Dugoujon Jean-Michel, Fortes-Lima Cesar, Migot-Nabias Florence, Massougbodji Achille, Bellis Gil, Pereira Luisa, Gísli Másson Augustine Kong, Kári Stefánsson, and Agnar Helgason. Reconstructing an African haploid genome from the 18th century. Nature Genetics, 50(2):199–205, Feb 2018.

[35] Wertenbroek R, Hofmeister RJ, Xenarios I, Thoma Y, Delaneau O (2024) Improving population scale statistical phasing with whole-genome sequencing data. PLoS Genet 20(7): e1011092.

[36] Loh PR, Palamara PF, Price AL. Fast and accurate long-range phasing in a UK Biobank cohort. Nat Genet. 2016 Jul;48(7):811–6.

[37] James P Cook, Anubha Mahajan, Andrew P Morris, Fine-scale population structure in the UK Biobank: implications for genome-wide association studies, Human Molecular Genetics, Volume 29, Issue 16, 15 August 2020, Pages 2803–2811

[38] Monica D. Ramstetter, Sushila A. Shenoy, Thomas D. Dyer, Donna M. Lehman, Joanne E. Curran, Ravindranath Duggirala, John Blangero, Jason G. Mezey, Amy L. Williams, Inferring Identical-by-Descent Sharing of Sample Ancestors Promotes High-Resolution Relative Detection, The American Journal of Human Genetics, Volume 103, Issue 1, 2018, Pages 30–44

[39] Michael P. Epstein, William L. Duren, and Michael Boehnke, Improved Inference of Relationship for Pairs of Individuals, Am. J. Hum. Genet. 67:1219–1231, 2000.

[40] Tahmasbi R, Keller MC. GeneEvolve: a fast and memory efficient forward-time simulator of realistic whole-genome sequence and SNP data. Bioinformatics. 2017 Jan 15;33(2):294–296.

[41] Halldorsson BV, Palsson G, Stefansson OA, Jonsson H, Hardarson MT, Eggertsson HP, Gunnarsson B, Oddsson A, Halldorsson GH, Zink F, Gudjonsson SA, Frigge ML, Thorleifsson G, Sigurdsson A, Stacey SN, Sulem P, Masson G, Helgason A, Gudbjartsson DF, Thorsteinsdottir U, Stefansson K. Characterizing mutagenic effects of recombination through a sequence-level genetic map. Science. 2019 Jan 25;363(6425).

