## Supplement for "A Novel Method for Across-Chromosome Phasing without Relative Data" (from another version of this preprint: version 1, 2026-02-17)

### 1 Correction of within-chromosome phasing

The accuracy of across-chromosomal phasing depends heavily on the accuracy of within-chromosome phasing. UK Biobank has released data phased using Shapeit4, but we detected some phase errors (switching of haplotypes) in this data. Thus our first step was to correct the within-chromosome phasing in order to provide appropriate input for later steps. This correction was applied to all individuals in the sample to reduce the number of genotyping errors and phasing errors in the input data. The offspring of 978 parent/offspring trios and 3,718 parent/offspring duos were used for ground truth comparisons to assess the accuracy of the corrections. In order to make our corrections equally accurate for offspring and the rest of the sample, each person is corrected using only other individuals in the sample with whom they have a $\hat{\pi}$ lower than 0.1875.

#### 1.1 Correction based on number of segments

For each individual, we process one SNP at a time (including homozygous SNPs, which can still be affected by genotyping errors but not phase errors). We begin by identifying other individuals who have one haplotype that perfectly matches the focal individual's haplotype to the left or right of (eg. preceding or following) the focal SNP. We chose a minimum matching segment length of 128 SNPs (2 $\times$ 64 SNPs on Figure 1) (aproximately 1.2 centimorgans) to balance the need to collect sufficient comparison segments against the need to minimize the number of random matches (matches that are not due to distant relatedness). Using a segment length which was a multiple of 64 bits also allowed for a more computationally efficient implementation of the comparison (on a 64-bit machine).

For each of the focal individual's SNPs, we calculate eight weights: $WRA(0)$, $WRA(1)$, $WLA(0)$, $WLA(1)$, $WRB(0)$, $WRB(1)$, $WLB(0)$, $WLB(1)$. The letters $L$ and $R$ indicate whether the matching segment is to the left or right of the focal SNP, the letters $A$ and $B$ indicate which of the focal individual's haplotypes is being compared, and the numbers 0 or 1 indicate genotype at the focal SNP. For example, weights beginning with $WLB$ are based on haplotypes from other individuals which match the focal individual's haplotype B for the segment to the left of the focal SNP, with $WLB(1)$ representing haplotypes where the focal SNP has genotype 1 and $WLB(0)$ representing haplotypes where the focal SNP has genotype 0.

For each of the four potential matching segments there is a pair of weights corresponding to the two potential genotypes (eg. $WLA(0)$ and $WLA(1)$ for the segment to the left of focal SNP on the focal individual's haplotype A). These paired weights are compared to each-other to assign values of "0" "1" or "inconclusive" to the four variables $LA$ $LB$ $RA$ and $RB$ (for example, if $WLA(1) > 2 \times WLA(0)$ then $LA = 1$ otherwise $LA = inconclusive$). These four values are then compared to focal individual's two haplotypic genotypes for the focal SNP ($FA$ and $FB$), with inconsistencies indicating potential errors in the genetic data.

The Figure 1 describes this process. The two thresholds $SW$ and $PW$ determine if the difference between the genotype-0 and genotype-1 weights for a SNP should be taken into account (if the difference is not taken into account, the corresponding variable is set to "inconclusive"). The four variables $LA$, $LB$, $RA$, and $RB$ can each take three different values (0, 1, inconclusive) which leads to 81 (3 $\times$ 3 $\times$ 3 $\times$ 3) possible combinations. Among these combinations, those where $LA$ and $LB$ (or $RA$ and $RB$) are inconsistent with the focal SNP's genotypes on each haplotype may be the result of phase errors or genotyping errors.

Any scenario where at least two of the four variables $LA$, $LB$, $RA$, and $RB$ are inconsistent with an individual's reported focal SNP genotypes ($FA$ and $FB$) is evaluated as a potential error. Genotyping errors and phase errors can be distinguished from other situations such as meiotic recombination or deletions/duplications by a threshold of 2/4 mismatches. In the case of recombination, only one of an individual's two haplotypic genotypes for the focal SNP will differ from those predicted by matching individuals, and only on one side of the focal SNP (for example, $FA$ may differ from either $LA$ or $RA$). Deletions and duplications are likely to lead to some/all of the variables $LA$, $LB$, $RA$, or $RB$ being assigned the value "inconclusive" due to a lack of matching segments.

For each possible combination of at least 2/4 mismatches (eg. "$LA$ and $LB$ match phased focal SNP genotypes, $RA$ and $RB$ do not match"), we use ground-truth phasing information from the 978 trio offspring and the 3718 parent-offspring duos to set thresholds for $SW$ and $PW$ that will most accurately distinguish errors from correct genotypes for that type of mismatch. These thresholds are then applied for all individuals in the sample and used to correct all the SNPs for which each combination applies. Errors detected during this step may be corrected by reversing the phase of the chromosome from the focal SNP until to address a phase error the end or by changing a genotype to address a genotyping error. This process is repeated for five iterations, so the errors that are corrected after one run can help to detect new errors in the following runs. These initial corrections address genotyping errors and some phase errors.

Following the initial corrections, we perform additional detection and correction of phase errors. For this step, we focus on individuals who match the focal individual for at least 64 SNPs on both sides of the focal SNP (unlike the previous step which required only one side to match), and check whether the left and right matching regions both match to the same haplotype of the focal individual. This produces two sets of individuals: $same\_hap$ (whose left-side and right-side matches both match to the same haplotype of the focal individual) and $diff\_hap$ (whose left-side and right-side matches are for opposite haplotypes of the focal individual). For each side (left or right), each haplotype (A or B), and each set of matching individuals ($same$ or $diff$), we calculate a weight using the same formula as $W_h$ in Figure 1, using 64 SNPs rather than 128. This produces eight weights, $WRA(same)$, $WRA(diff)$, $WLA(same)$, $WLA(diff)$, $WRB(same)$, $WRB(diff)$, $WLB(same)$, $WLB(diff)$. For each pair of two weights, we use ground truth data from the parent-offspring trios and duos to establish thresholds for $SW$ and $PW$ that will identify phase errors. If the selected values for these thresholds would improve the phasing in offspring when compared to ground truth, corrections using these two thresholds are applied to the entire sample. Errors detected during this step are corrected by reversing the phase of all of the focal individual's heterogzygous SNPs to the right of the focal SNP.

### 2 Algorithm

### 3 Composition of the sample

Figure 2 shows the distribution of $\hat{\pi}$ values in a sample of 17,000 focal individuals. For each individual, their highest $\hat{\pi}$ with another individual in the sample is shown, excluding close relatives ($\hat{\pi}$ > 0.33). The number of individuals whose closest relative is of degree 2, 3, 4, 5, or more distant can be deduced from the $\hat{\pi}$ values. Table 1 compares the percentage of focal individuals in the trio offspring and in a sample of 17,000 individuals having a closest relative of degree 2, 3 , 4, 5 or more according to $\hat{\pi}$.

**Algorithm 1:** Algorithm for detection errors

**input** : The genome of individuals for a chromosome
**parameter:** $SumWeight$,$ProductWeight$
**Result:** The four variables $LA$, $LB$, $RA$, and $RB$
**for** *$snp$ in chromosome* **do**
  **for** *Individual $I$* **do**
    **for** *haplotype $h$ in (A, B)* **do**
      **for** *$D$ in (Left, Right)* **do**
        Set the 2 counters $W0$ and $W1$ to 0 **for** *Unrelated $U$* **do**
          $k$ = number of consecutive perfect matches of 128 SNPs between $I$ and $U$ starting from SNP $snp$ in the direction $D$
          **if** *$k$ > 0* **then**
            $Wh = Wh$ + (k - 1) × k × (2 × k + 1)
        **if** *$W0$+$W1$>$SumW$ **and** max($W0$,$W1$)>min($W0$,$W1$) × $ProductW$* **then**
          **if** *$W0 > W1$* **then**
            $Dhap$=0
          **else**
            $Dhap$=1
        **else**
          $Dhap$=inconclusive

**Algorithm 2:** Main Algorithm

**Result:** across Phasing of an individual
Following algorithm 1, $W_1$ is determined
Meta chromosome ← $W_1$
**for** *$w$ in {remaining window}* **do**
  Following algorithm 1: $W_2$ ← merging chromosome
  Meta chromosome ← merge(Meta chromosome, $W_2$)

| | Degree 2 | Degree 3 | Degree 4 | Degree 5 | More Distant |
|---|---|---|---|---|---|
| 978 Trio Offspring | 16.0% | 10.6% | 30.5% | 25.4% | 17.5% |
| Random 17000 Individuals | 4.1% | 18.9% | 20.0% | 32.6% | 24.4% |

Table 1. Comparison of trio offspring and random individuals by the degree of their closest non-first-degree relative in the sample. For offspring of trio data and random individuals in the dataset percentage of individuals with a closest relative (excluding degree 1) of degree 2, 3, 4, 5 or more according to $\hat{\pi}$.

| | | | Degree 2 | Degree 3 | Degree 4 | Degree 5 | More Distant | ALL |
|---|---|---|---|---|---|---|---|---|
| Shapeit2 Phasing | Our method | Trio offspring | 94.45 | 91.06 | 85.54 | 81.16 | 72.35 | 85.24 |
| | | P/O offspring | 91.20 | 90.98 | 86.14 | 80.12 | 71.76 | 84.57 |
| | Noto's method | Trio offspring | 82.2 | 82 | 76.5 | 67.9 | 61.5 | 74.5 |
| | | P/O offspring | 80.4 | 75.8 | 71.1 | 61.4 | 56.4 | 67.8 |
| Parental Phasing | Our method | Trio offspring | 100 | 100 | 100 | 96.5 | 89.2 | 100 |
| | | P/O offspring | 100 | 100 | 98.2 | 90 | 86.6 | 96.6 |
| | Noto's method | Trio offspring | 100 | 100 | 96.9 | 92 | 80 | 96.5 |
| | | P/O offspring | 100 | 100 | 97.2 | 87.9 | 86.4 | 95.4 |
| | | | Degree 2 | Degree 3 | Degree 4 | Degree 5 | More Distant | ALL |
| Shapeit2 Phasing | Our method | Trio offspring | 93.92 | 89.66 | 83.96 | 78.21 | 70.84 | 82.40 |
| | | P/O offspring | 90.40 | 89.73 | 84.21 | 78.47 | 70.68 | 81.73 |
| | Noto's method | Trio offspring | 82 | 81.1 | 75 | 69.8 | 62.6 | 73.3 |
| | | P/O offspring | 79.9 | 75.6 | 70.6 | 62.8 | 57.8 | 68.1 |
| Parental Phasing | Our method | Trio offspring | 99.9 | 99.5 | 97.7 | 93.9 | 86.6 | 95.3 |
| | | P/O offspring | 99.8 | 99.2 | 96.4 | 87.9 | 71 | 90.6 |
| | Noto's method | Trio offspring | 99.4 | 98.2 | 95.1 | 89.2 | 76.9 | 91.4 |
| | | P/O offspring | 99.7 | 99.1 | 96 | 85.4 | 69.8 | 89.6 |

Table 2. Median (top) and Mean (bottom) Scores of our method vs. Noto et. al's IBD-based method. Results are compared according to how the offspring were phased (Shapeit2 or parental phasing) and whether the offspring were from parent-child trios or from duos with only one parent.

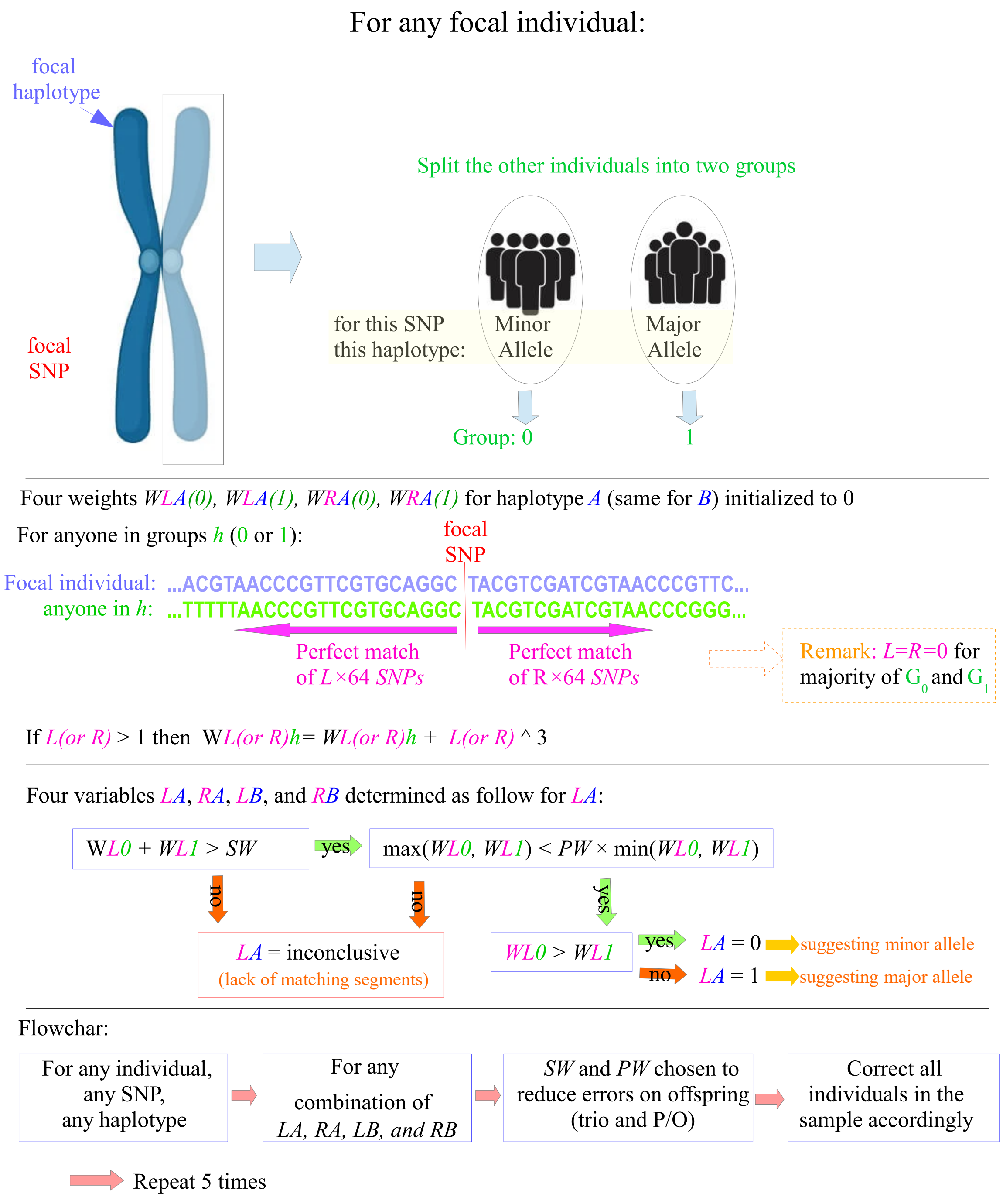


Figure 1: Schema for detection of phase and genotype errors

#### 4 Heatmap of $\lambda$ values

The positive values of $\lambda$ observed along the diagonal predominantly correspond to pairs of genomic windows situated on the same chromosome and are attributable to the presence of identity-by-descent (IBD) segments.

#### 5 Number of samples

The sample size within the dataset (in this case, 435,187 individuals) constitutes a critical parameter influencing the performance of the algorithm, alongside the inclusion of relatives of the focal individual. The present analysis examines the effect of reduced sample sizes while maintaining a constant number of both close and distant relatives for the focal subject. For each focal individual, subsamples are constructed comprising the ten

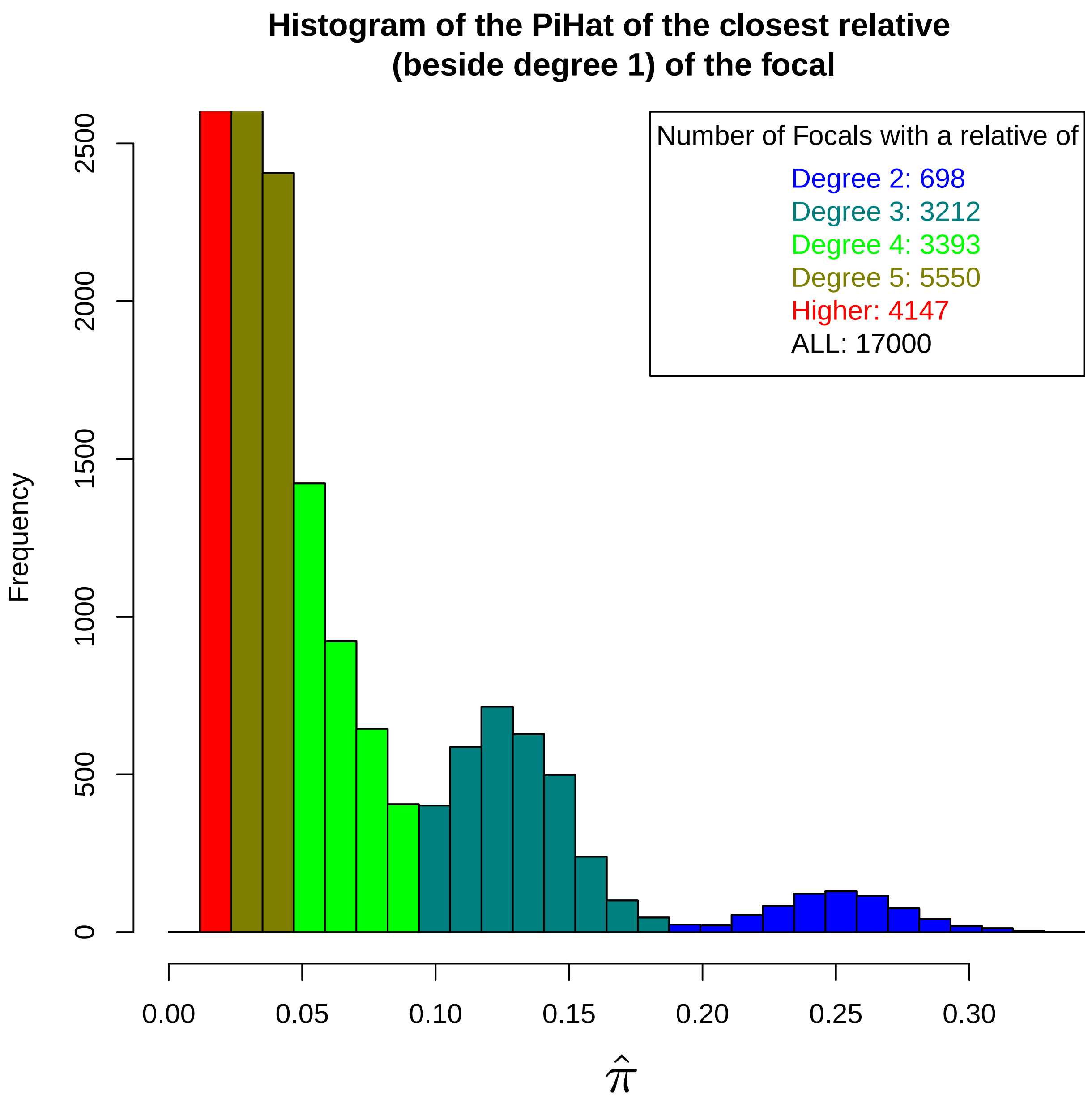


Figure 2: Distribution over 17,000 random individuals of the $\hat{\pi}$ of the closest relative of each individual (excluding degree 1 relative) in the sample.

closest relatives (as determined by the estimated $\hat{\pi}$) and additional individuals, with total subsample sizes defined as $\frac{1}{5^h}$ of the original dataset, where $h$ ranges from 1 to 5.

#### 6 Number of phase errors

The quality of within chromosome phasing has a direct impact on across chromosome phasing. To examine this, we generated datatsets with a varying proportion of within chromosome phase errors. We began by generating data with fewer phase errors by using the parental genomes to correct 1/3, 2/3, or all within-chromosome phase errors in offspring. We also generated data with more phase errors by adding either 2x or 5x as many phase errors to each offspring's data. Phase errors are not randomly located as they are more frequent near recombination hotspots. In order to have realistic phase errors, for every offspring, we added phase errors located on a SNP on which a identified phase error occurred in another offspring.

The results, presented Figure 6, show the across chromosome phasing is more robust for offspring with relatives. Close relative may allow the

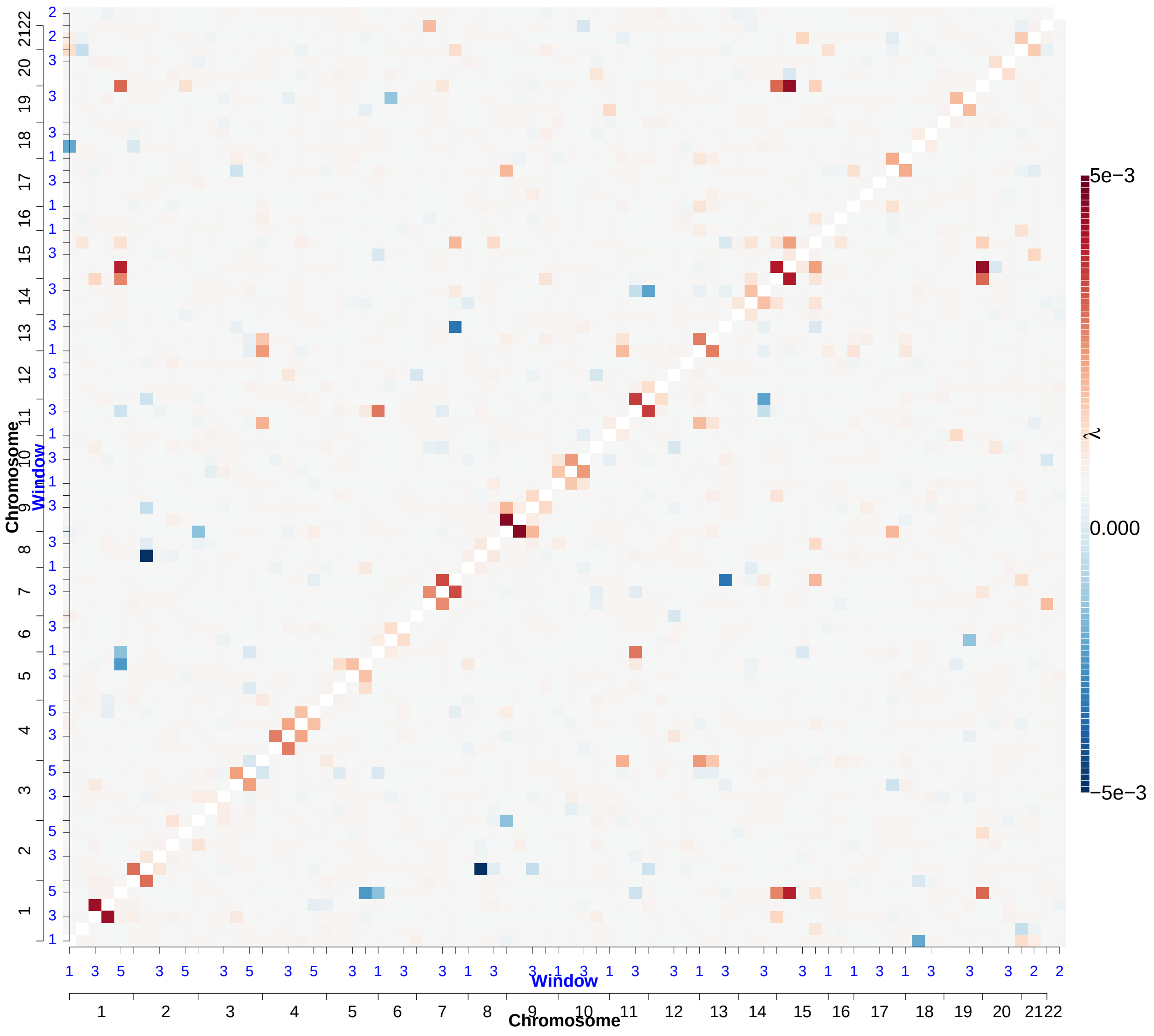


Figure 3: Heat map of $\lambda$ values for an individual at the first iteration.

algorithm to phase accurately two windows on the two sides of a phase error.

#### 7 Visualization of Phasing Evolution

To facilitate the interpretation of the iterative merging process, we developed a visualization framework that captures the evolution of the across-chromosome phase. This is achieved through a sequence of graphical representations where each frame represents a single iteration of the merging algorithm.

The visualization integrates three primary components:

- **Merging Map:** A genome-wide representation of the chromosomal windows. Windows that remain unmerged are represented in grey. Once a window is incorporated into the growing meta-chromosome, it is assigned a specific color corresponding to its assigned group. This allows for the observation of the algorithm's progression as it jumps between chromosomes based on the maximum absolute value of $\lambda$.
- **Dynamic Heatmap:** A real-time update of the $\lambda$ correlation matrix. As the algorithm iterates, the heatmap illustrates the transition from a global matrix of unphased windows to an organized state, highlighting

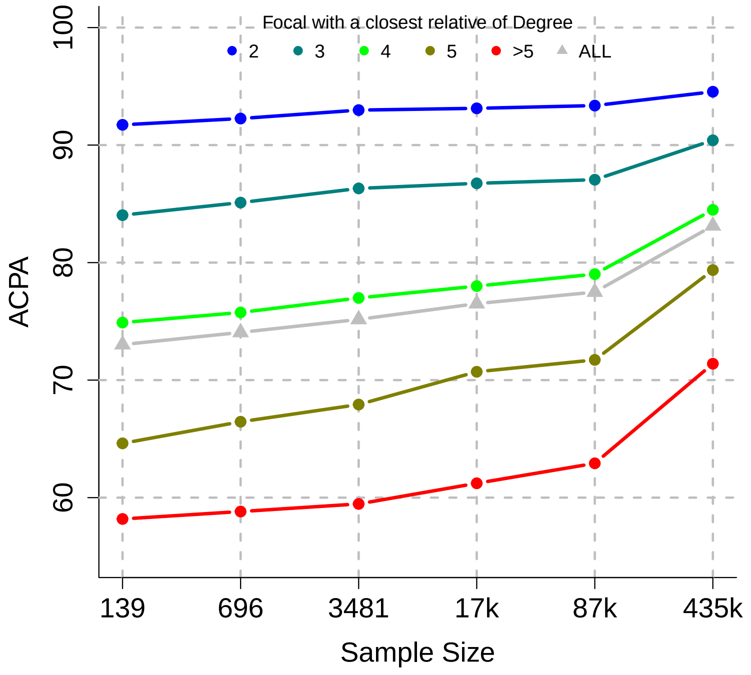


Figure 4: Mean ACPA for offspring of trio data with different sample sizes.

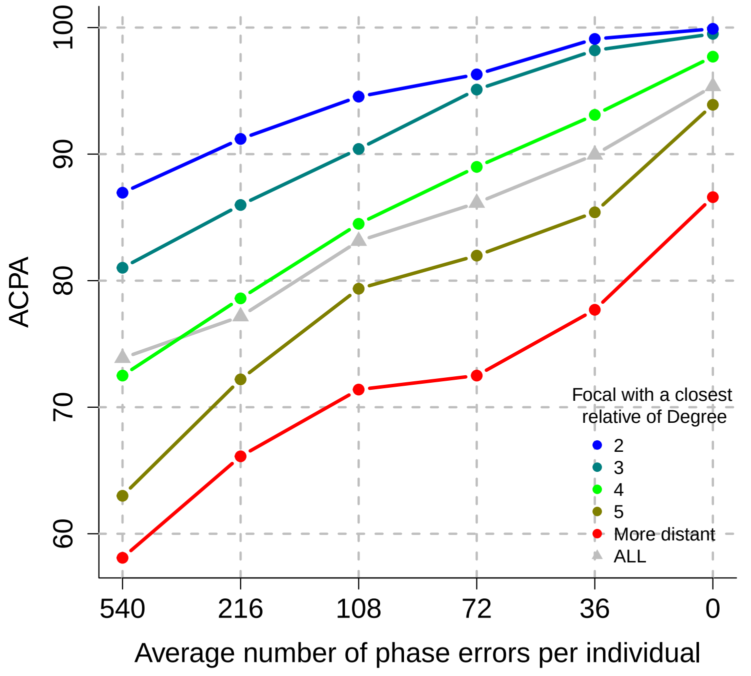


Figure 5: Mean ACPA for offspring of trio data with different rate of phase errors.

how noise is reduced as more windows are correctly aligned to the focal meta-chromosome.

- **Distribution Histograms:** A live histogram tracking the $\lambda$ values for all remaining window pairs. This diagnostic tool monitors the "signal-to-noise" ratio during execution. As the algorithm progresses, a clear bimodal distribution typically emerges, providing a statistical signature that separates pairs inherited from the same parent from those inherited from different parents.

This visual approach provides a qualitative diagnostic tool to identify regions where within-chromosome phase errors may be causing conflicting $\lambda$ signals, which appear as inconsistent merging patterns in the observed sequence.

#### 8 Windows

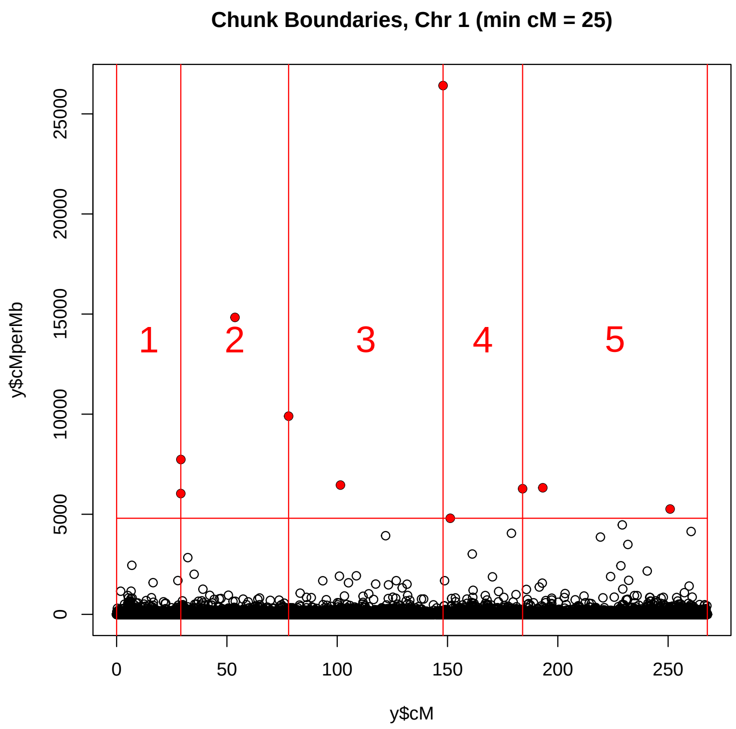


Figure 6: Map of window boundaries on chromosome 1.

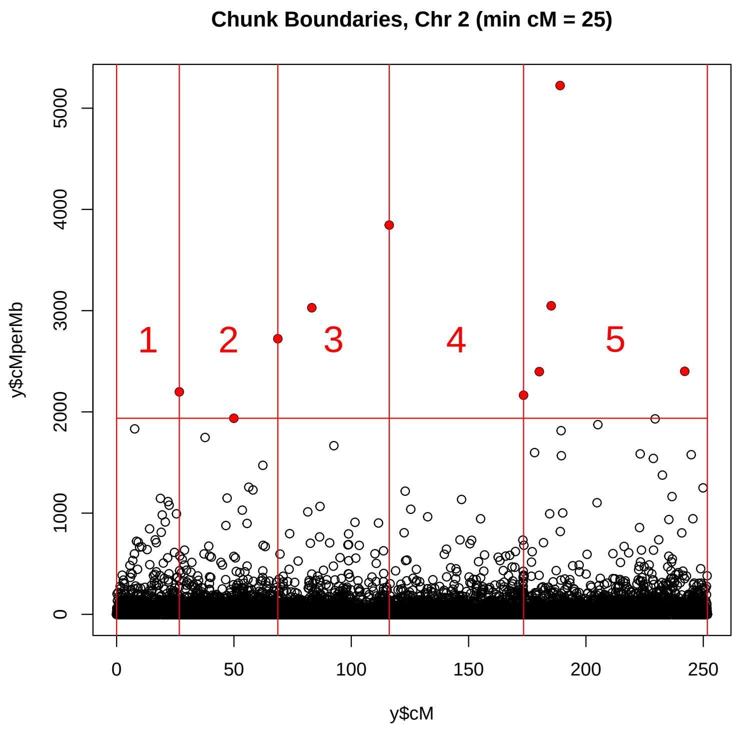


Figure 7: Map of window boundaries on chromosome 2.

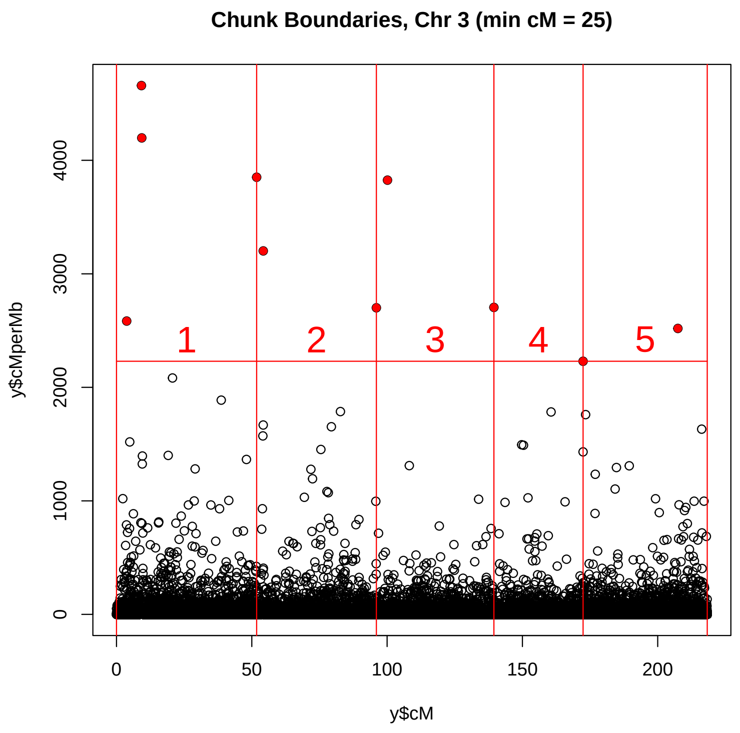


Figure 8: Map of window boundaries on chromosome 3.

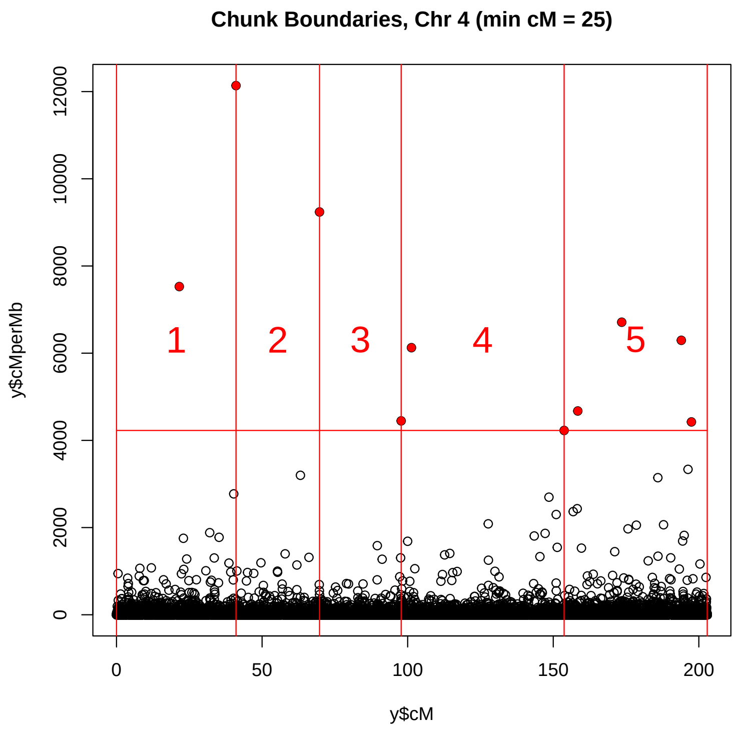


Figure 9: Map of window boundaries on chromosome 4.

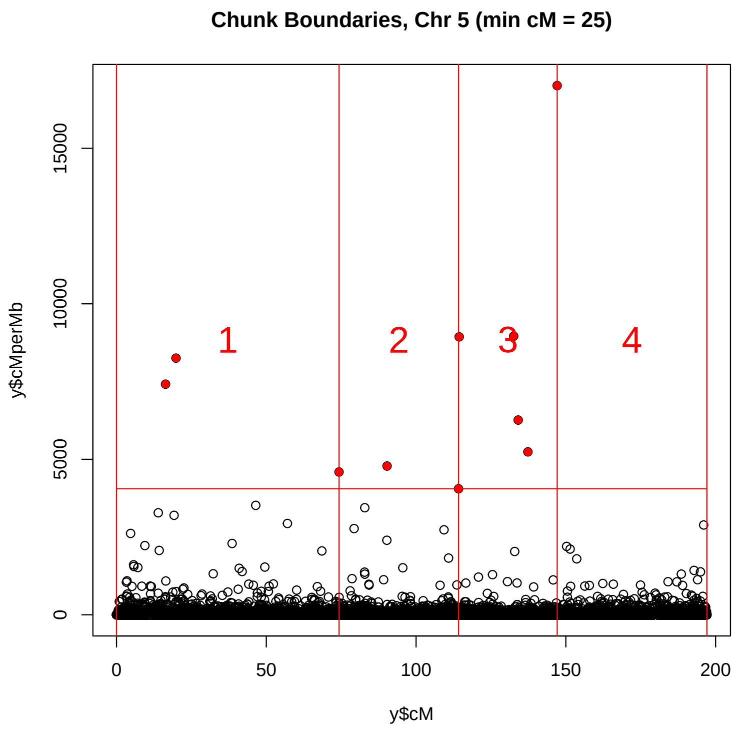


Figure 10: Map of window boundaries on chromosome 5.

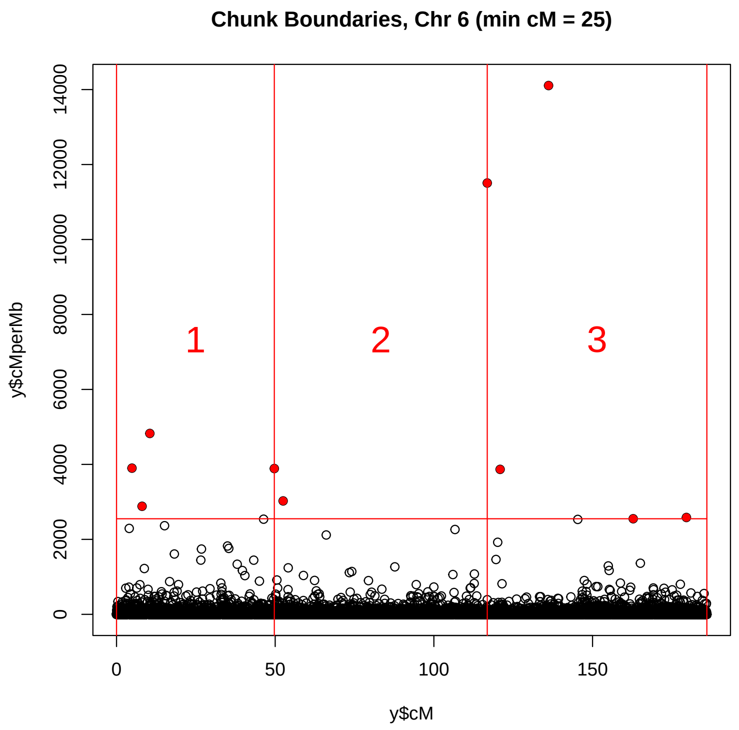


Figure 11: Map of window boundaries on chromosome 6.

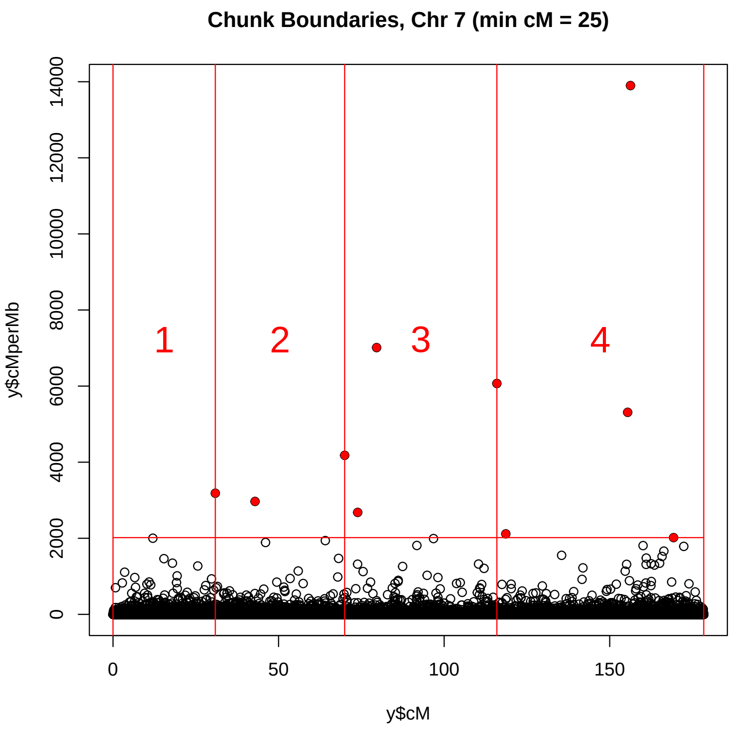


Figure 12: Map of window boundaries on chromosome 7.

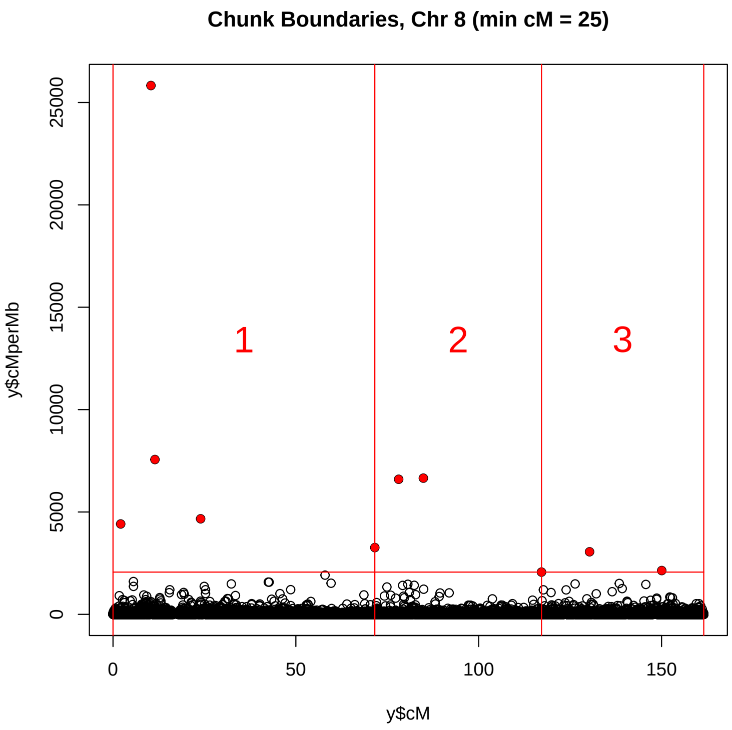


Figure 13: Map of window boundaries on chromosome 8.

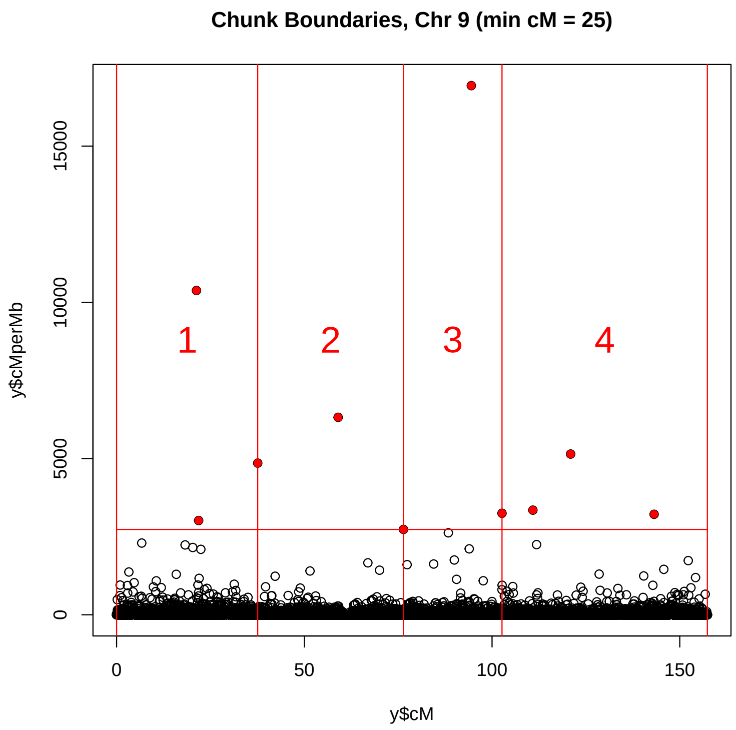


Figure 14: Map of window boundaries on chromosome 9.

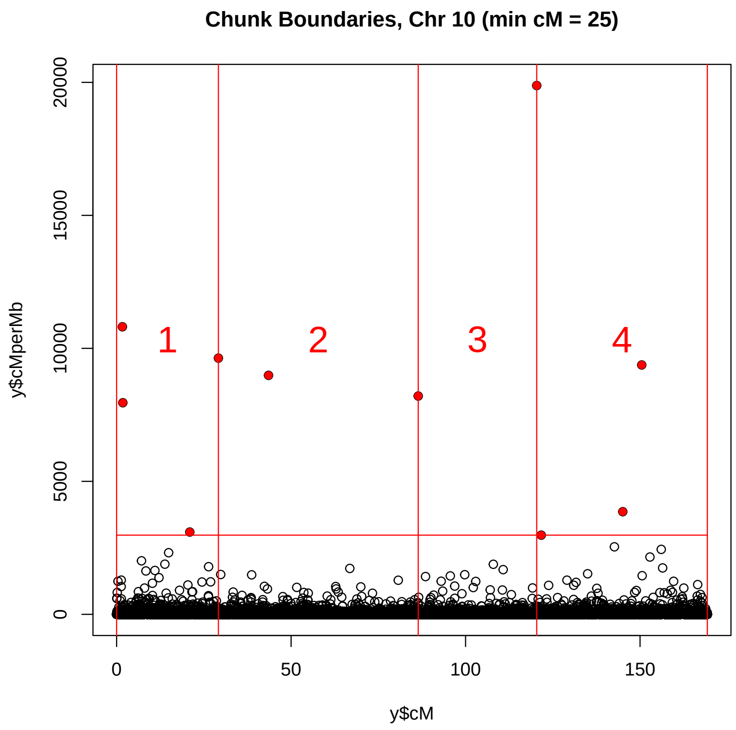


Figure 15: Map of window boundaries on chromosome 10.

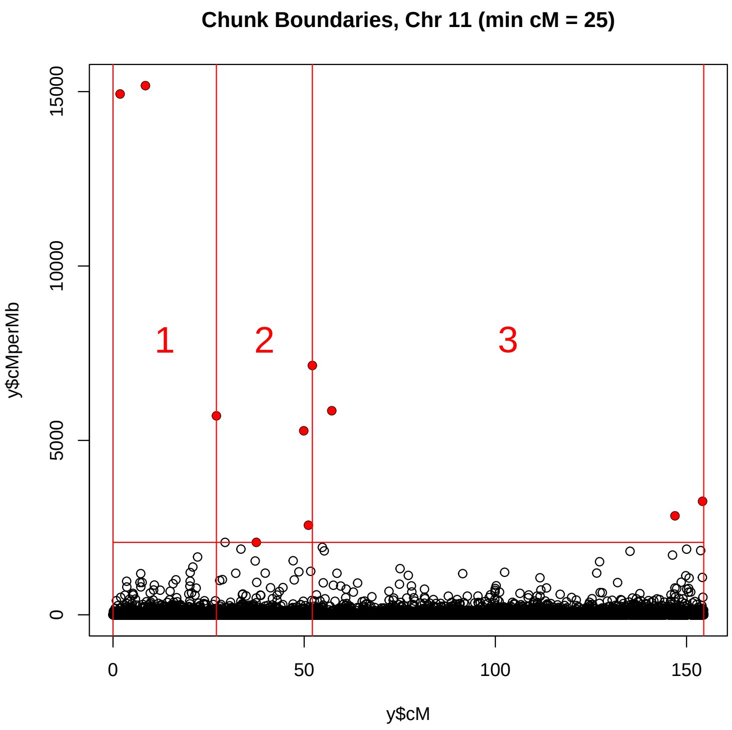


Figure 16: Map of window boundaries on chromosome 11.

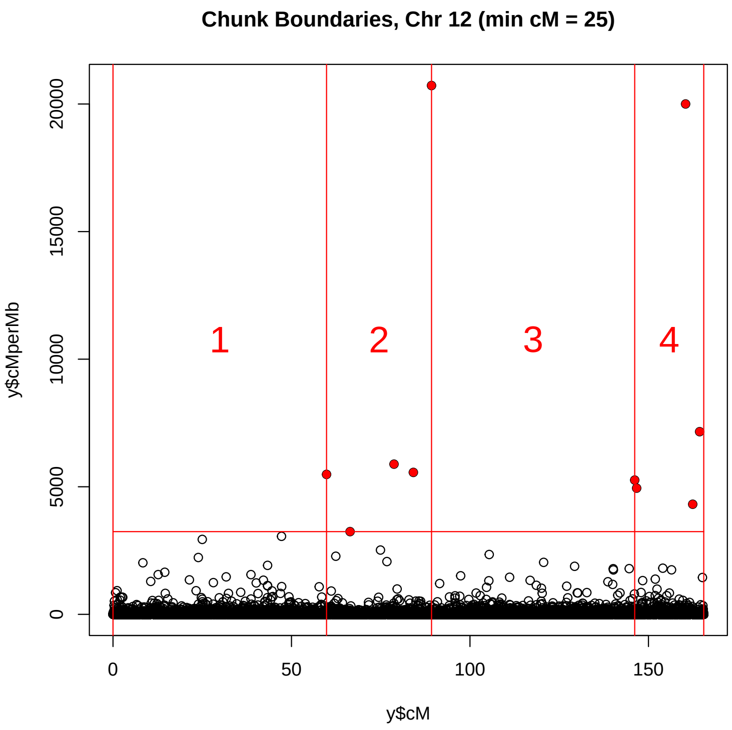


Figure 17: Map of window boundaries on chromosome 12.

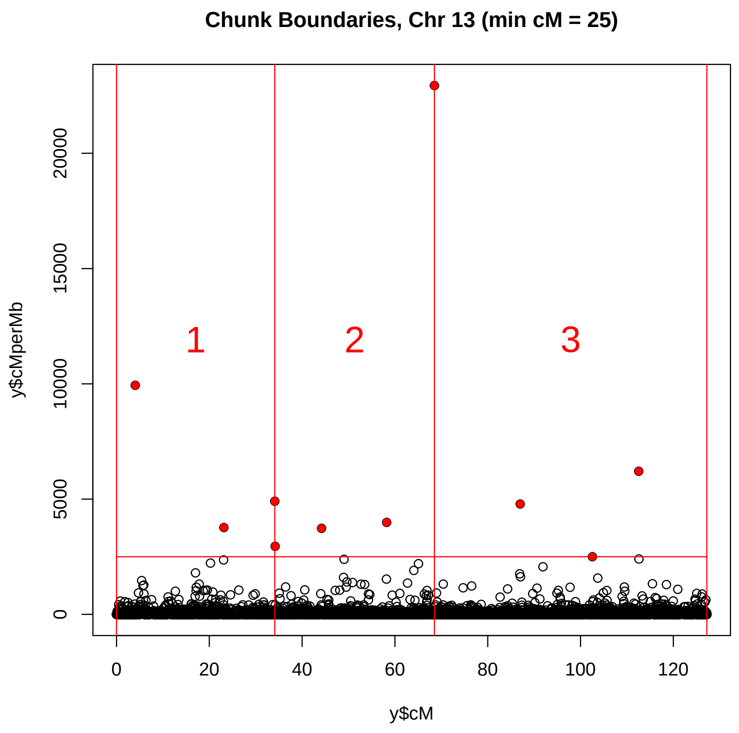


Figure 18: Map of window boundaries on chromosome 13.

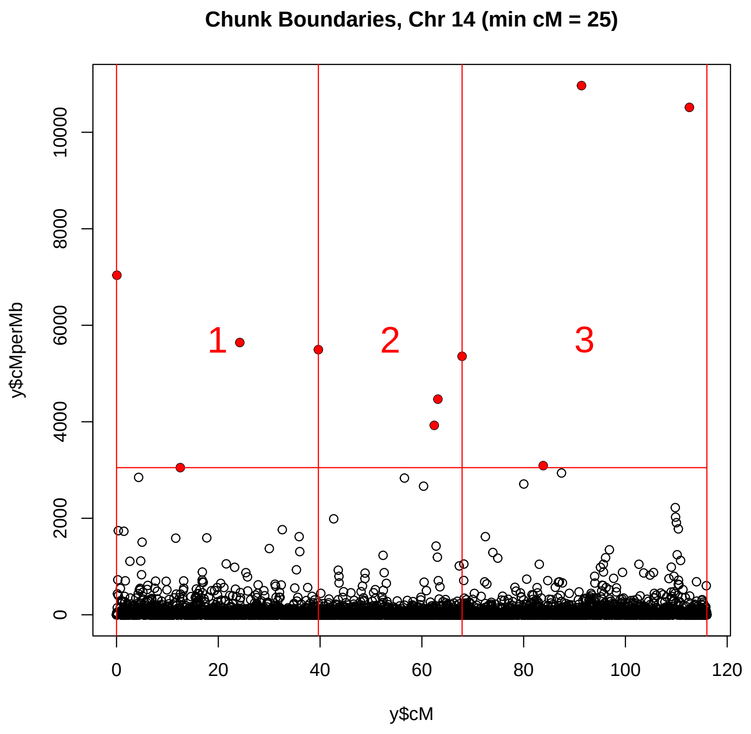


Figure 19: Map of window boundaries on chromosome 14.

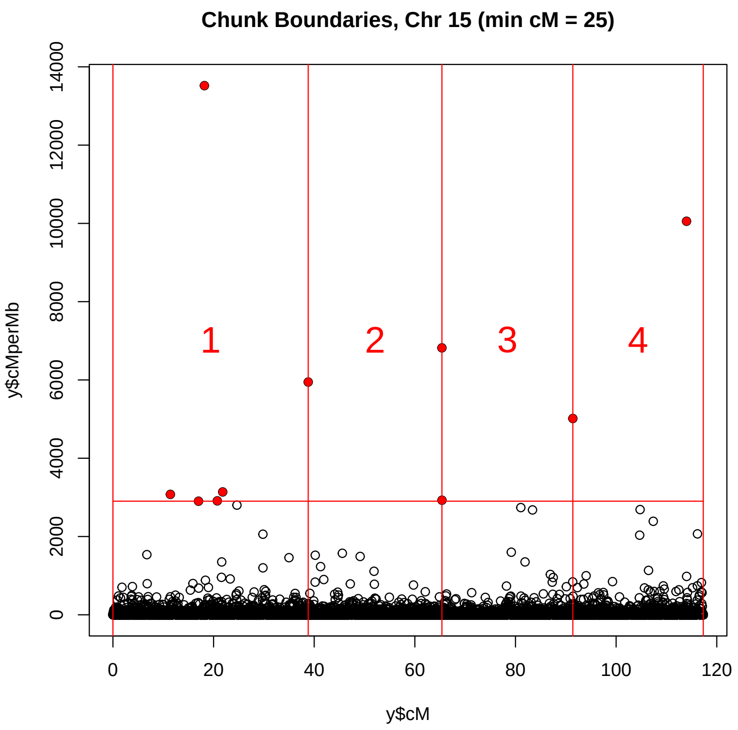


Figure 20: Map of window boundaries on chromosome 15.

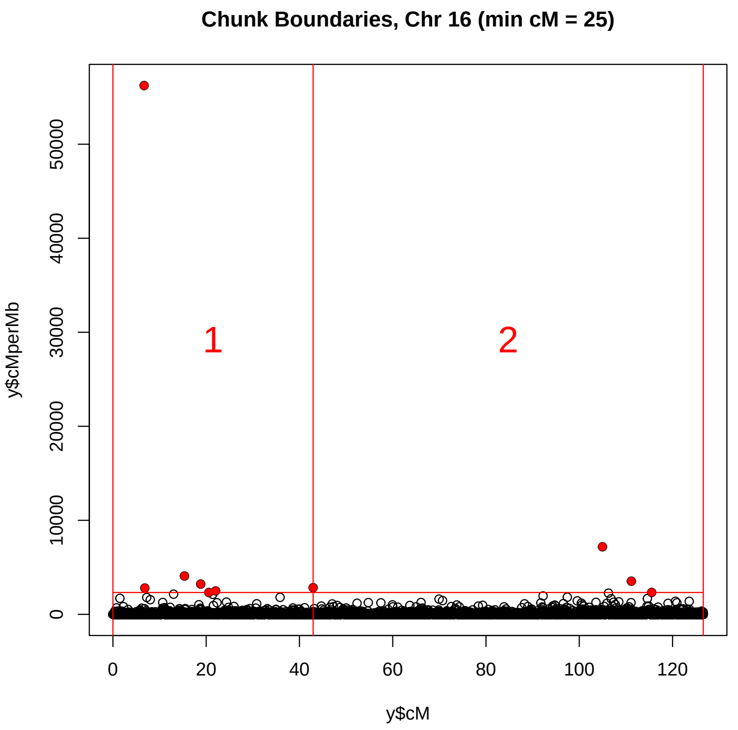


Figure 21: Map of window boundaries on chromosome 16.

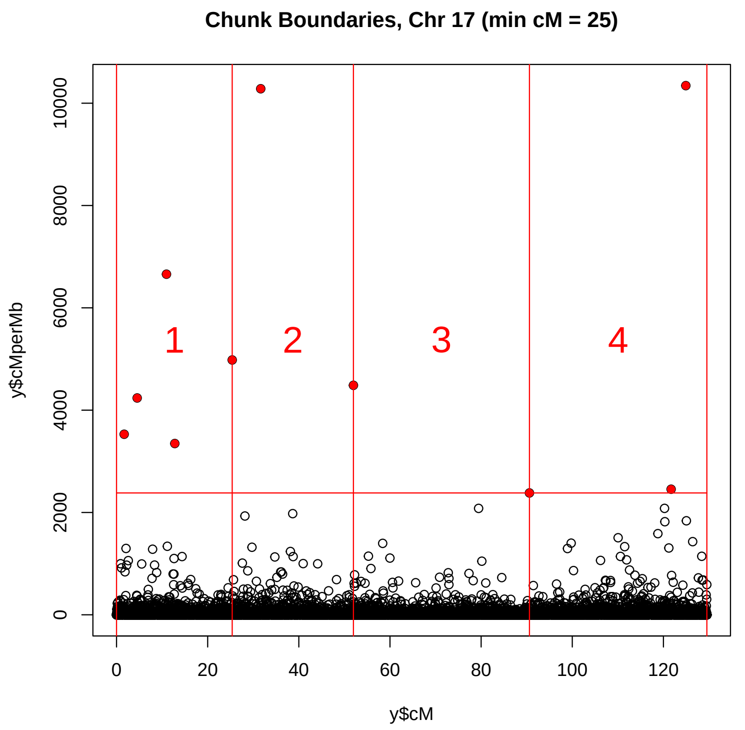


Figure 22: Map of window boundaries on chromosome 17.

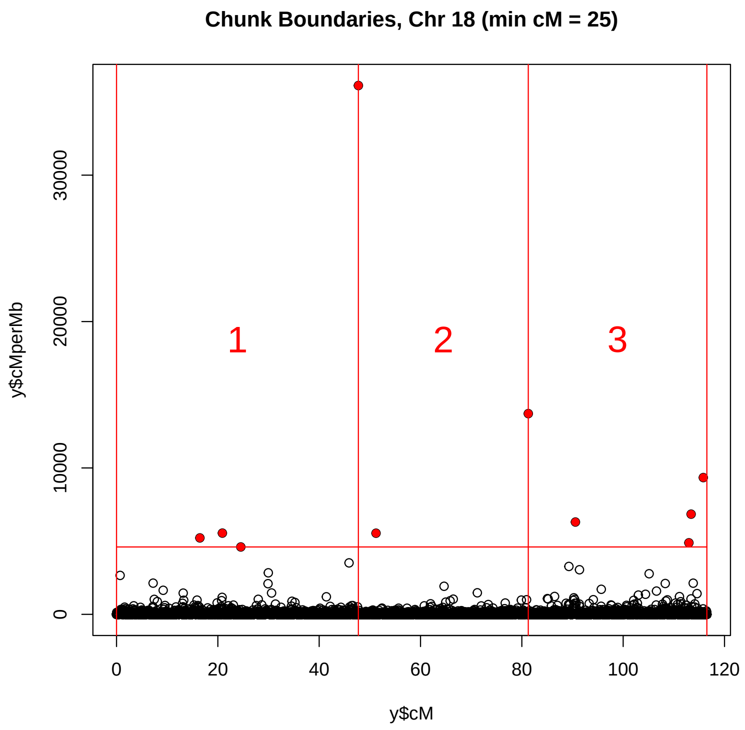


Figure 23: Map of window boundaries on chromosome 18.

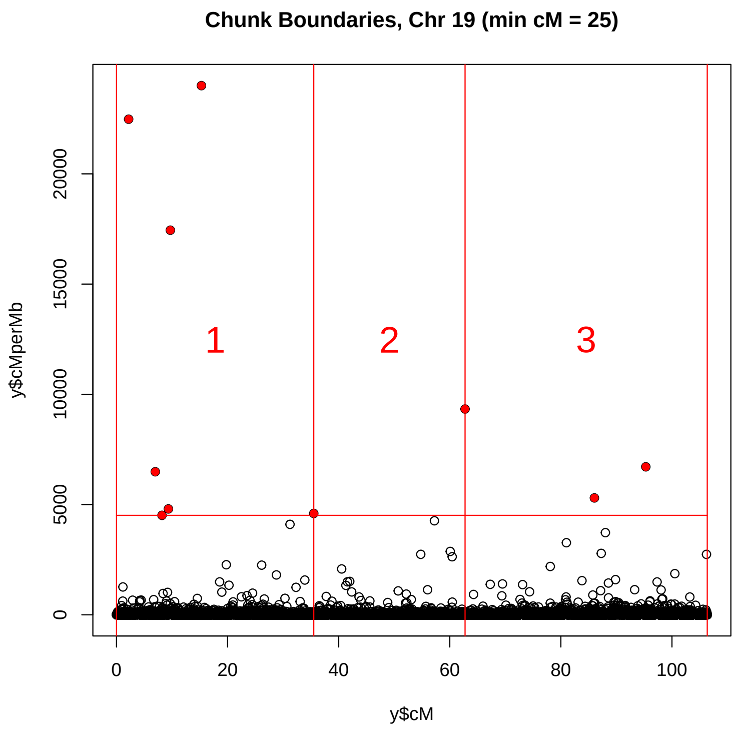


Figure 24: Map of window boundaries on chromosome 19.

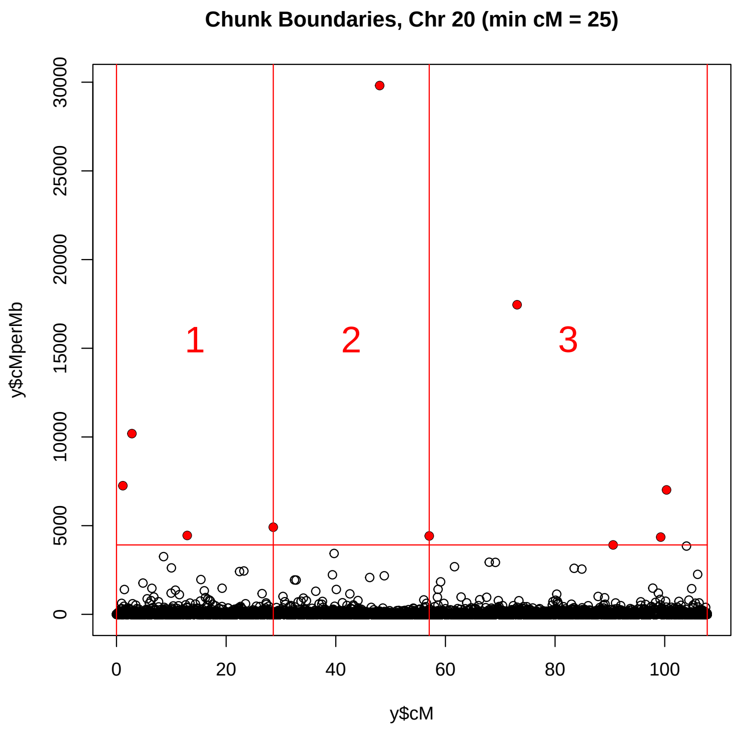


Figure 25: Map of window boundaries on chromosome 20.

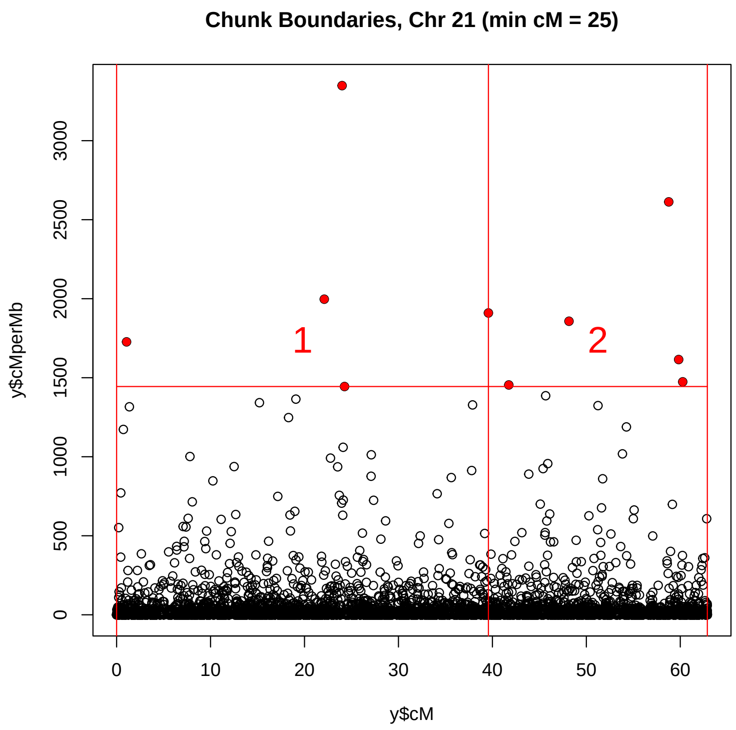


Figure 26: Map of window boundaries on chromosome 21.

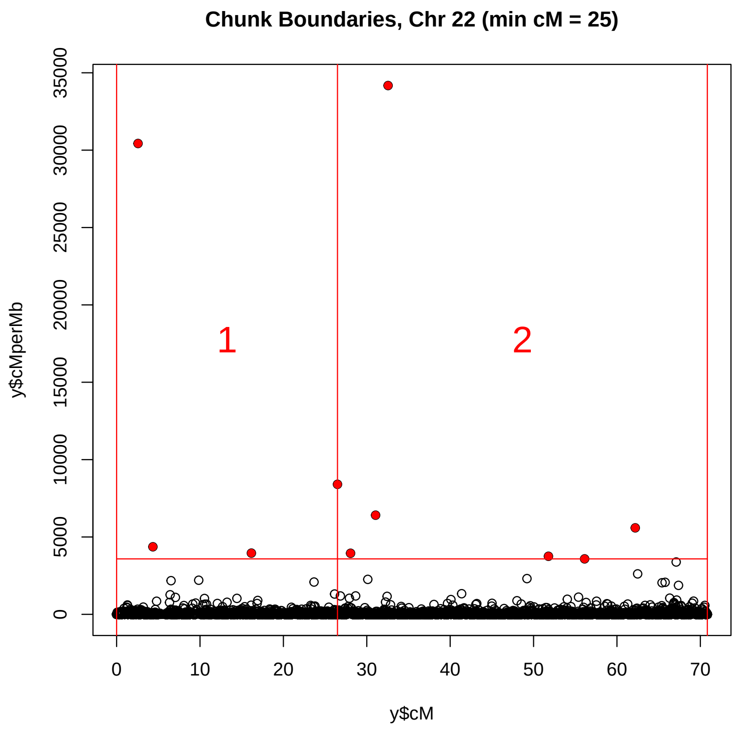


Figure 27: Map of window boundaries on chromosome 22.
